# Midbrain dopamine D2R regulates the salience of threat-related events

**DOI:** 10.1101/2024.11.12.623233

**Authors:** Laia Castell, Angelina Rogliardo, Tinaig Le Borgne, Claire Naon, Adeline Cathala, Emma Puighermanal, Leila Makrini, Nejmeh Mashhour, Chloé P. Petit, Claire Bernat, Régis Nouvian, Laura Cutando, Jean-Michel Revest, Philippe Faure, Antoine Besnard, Camilla Bellone, Arnaud Monteil, Giuseppe Gangarossa, Fabio Marti, Emmanuel Valjent

**Affiliations:** IGF, Univ. Montpellier, CNRS, Inserm, F-34094 Montpellier, France; Plasticité du Cerveau CNRS UMR8249, École supérieure de physique et de chimie industrielles de la Ville de Paris (ESPCI), Paris, France; Univ. Bordeaux, INSERM, Neurocentre Magendie, U1215, F-33000 Bordeaux, France; Université Paris Cité, CNRS, Unité de Biologie Fonctionnelle et Adaptative, 75013 Paris, France; INM, Université. Montpellier, Inserm, F-34000 Montpellier, France; Department of Basic Neuroscience, University of Geneva, 1211, Geneva, Switzerland; Institut Universitaire de France (IUF), Paris, France; Department of Physiology, Faculty of Medicine Siriraj Hospital, Mahidol University, Bangkok, Thailand

**Keywords:** dopamine, D2 autoreceptor, salience, defensive behaviors

## Abstract

Salience attributed to stimuli predicting rewarding or aversive outcomes is critical for adaptive behavior. Dopamine (DA)-neurons play a central role in this process by modulating responses to both rewarding and aversive cues. DA-neurons are tightly and readily modulated by DA D2 autoreceptors (autoD2Rs), but their role in regulating responses to aversive stimuli remains unclear. In this study, we investigated the role of autoD2R in regulating the activity of VTA DA-neurons in response to salient aversive stimuli. Using *Drd2^Slc6a3^* mice, in which *Drd2* is selectively deleted in DA-neurons, we observed enhanced excitatory and inhibitory responses of VTA DA-neurons to aversive stimuli, suggesting that autoD2R acts as a critical regulatory brake. Importantly, this modulation occurred independently of either the pacemaker activity of DA-neurons or their coupling to the non-selective sodium leak channel NALCN. Behaviorally, *Drd2^Slc6a3^* male mice showed enhanced discrimination between threat-predicting and non-predicting cues that persisted during extinction learning, highlighting a sex-biased role of autoD2R in threat processing. Our results provide new mechanistic insights through which autoD2R influence behavioral responses to aversive stimuli, with implications for understanding neuropsychiatric disorders characterized by maladaptive threat processing.

## Introduction

The ability to attribute motivational salience to stimuli that predict rewarding or aversive outcomes is essential for survival. This process relies on associative learning, which enables salient cues to recruit attentional resources and assists animals to prioritize actions in response to significant events (Bromberg-Martin et al., 2010). However, the aberrant attribution of motivational salience to reward- or aversion-predicting cues can result in over- or under-responsiveness, often linked to impaired motivated behavior (Neumann & Linscott, 2018; Roiser et al., 2013). These alterations are frequently associated with core symptoms of neuropsychiatric disorders, such as generalized anxiety and high-risk taking behaviors.

Dopamine (DA)-neurons are a major component of the brain’s reward system (Lammel et al., 2014). While their role in coding reward prediction errors is well established (Schultz, 2016), DA-neurons also encode motivational salience of a stimulus regardless of its valence, whether positive or negative (Bromberg-Martin et al., 2010). Distinct midbrain DA-neurons play complementary roles in assigning motivational salience to reward- and/or aversive-related information and in optimizing the corresponding behaviors (Lin et al., 2021; Schultz, 2016).

Dopamine D2 receptors expressed by DA-neurons (autoD2Rs) exert a potent inhibitory feedback on midbrain DA transmission by regulating DA-neuron excitability as well as DA turnover, synthesis and release (Ford, 2014). Low levels of autoD2Rs in DA-neurons have been associated with increased locomotor activity in response to environmental changes and with heightened sensitivity to rewarding and motivational salience to psychostimulants-paired cues (Bello et al., 2011; Holroyd et al., 2015). Moreover, the pivotal role of autoD2Rs in the regulation of salience to reward-predicting cues is further highlighted by the strong correlation between low autoD2Rs availability and novelty-seeking traits in humans (Zald et al., 2008), which is a primary risk factors in the transition from casual to compulsive drug use (Howard et al., 1997). While autoD2Rs are essential for curbing impulsivity and reward-seeking behaviors, their role in regulating motivational salience to aversive cues predicting potential threats remains unexplored.

Here, we reveal the importance of D2R signaling in midbrain DA-neurons for regulating the activity of VTA DA-neurons in response to salient aversive stimuli. We demonstrate that these effects do not require the pacemaker activity of DA-neurons nor the coupling of autoD2Rs to the non-selective sodium leak channel NALCN. Finally, we found that in male mice autoD2Rs act as a brake to limit exacerbated defensive behaviors in response to threat-associated stimuli.

## Materials and methods

### Animals

C57Bl/6J mice (Charles River Laboratories, France) were used for *in situ* hybridization. *Drd2-Cre and Slc6a3-Cre* mice crossed with *Ribotag* mice were used for immunofluorescence analysis polysome immunoprecipitation. The deletion of *Drd2* and *Nalcn* from DA neurons was carried out by crossing *Slc6a3-Cre* mice with *Drd2^loxP/loxP^* (Bello et al., 2011) (*Drd2^Slc6a3^*) and *Nalcn^loxP/loxP^* mice (Flourakis et al., 2015) (*Nalcn^Slc6a3^*), respectively. Male and female mice *Drd2^Slc6a3^* and *Nalcn^Slc6a3^* were compared with controls (Cre-negative mice *Drd2^f/f^* and *Nalcn^f/f^*). Mice were housed in groups of 2 to 5 per cage (standard sizes according to the European animal welfare guidelines 2010/63/EU) and maintained in a 12h light/dark cycle (lights on from 7:00 am to 7:00 pm), in stable conditions of temperature (22°C) and humidity (60%), with food and water provided *ad libitum*. All animal procedures were conducted in accordance with the guidelines of the French Agriculture and Forestry Ministry for handling animals (authorization number/license B34-172-41) and approved by the relevant local and national ethics committees (authorizations APAFIS#38912 and APAFIS#22635).

### Histology

#### Immunofluorescence

Tissue preparation and immunofluorescence analyses were performed as previously described (Biever et al., 2015). Mice were anaesthetized with Euthasol® (360 mg/kg, i.p., TVM lab, France) and transcardially perfused with 4% (weight/vol) paraformaldehyde in 0.1 M sodium phosphate buffer (pH 7.5). Brains were post-fixed overnight in the same solution and stored at 4°C. Free-floating sections (40 µm) were washed in PBS, incubated 15 min in 0.2% Triton X-100 in PBS. Slices were then incubated in PBS 72 hrs at 4°C with the primary antibodies: rabbit anti-HA (1:1000, Rockland #600-401-384), rabbit mouse-TH (1:1000, Millipore #MAB318) and rat anti-DAT (1:1000, Millipore #MAB369). Slices were rinsed three times in PBS and incubated 45 min with goat Cy3-coupled anti-rabbit (1:500; Jackson ImmunoResearch Laboratories), goat Alexa Fluor 488-coupled anti-mouse and goat Alexa Fluor 647-coupled anti-rat (1:500; Jackson ImmunoResearch Laboratories) secondary antibodies. Sections were rinsed twice in PBS before mounting in 1,4-diazabicyclo-[2.2.2]-octane (DABCO, Sigma-Aldrich). Fluorescent images of labeled cells in the ventral tegmental area and *substantia nigra* were captured using sequential laser scanning confocal microscopy (Leica SP8). HA-positive cells were pseudocolored in green, while TH and DAT were pseudocolored in red. Images used for quantification of HA/TH-positive cells were all single confocal sections. Adjacent serial sections were never counted for the same marker to avoid any potential double counting of hemisected neurons. Values in the histograms in Figure 1 represent the estimated percentage of HA/TH co-expressing cells as percentage of TH-positive cells throughout the rostro-caudal extension of the rostro-caudal extension of the SNc and the VTA subareas including the parabrachial pigmented area (PBP), paranigral nucleus (PN), interfascicular nucleus (IF), and rostral linear nucleus (RLi) (4 levels per mouse, n = 3 mice).

**Figure 1:**
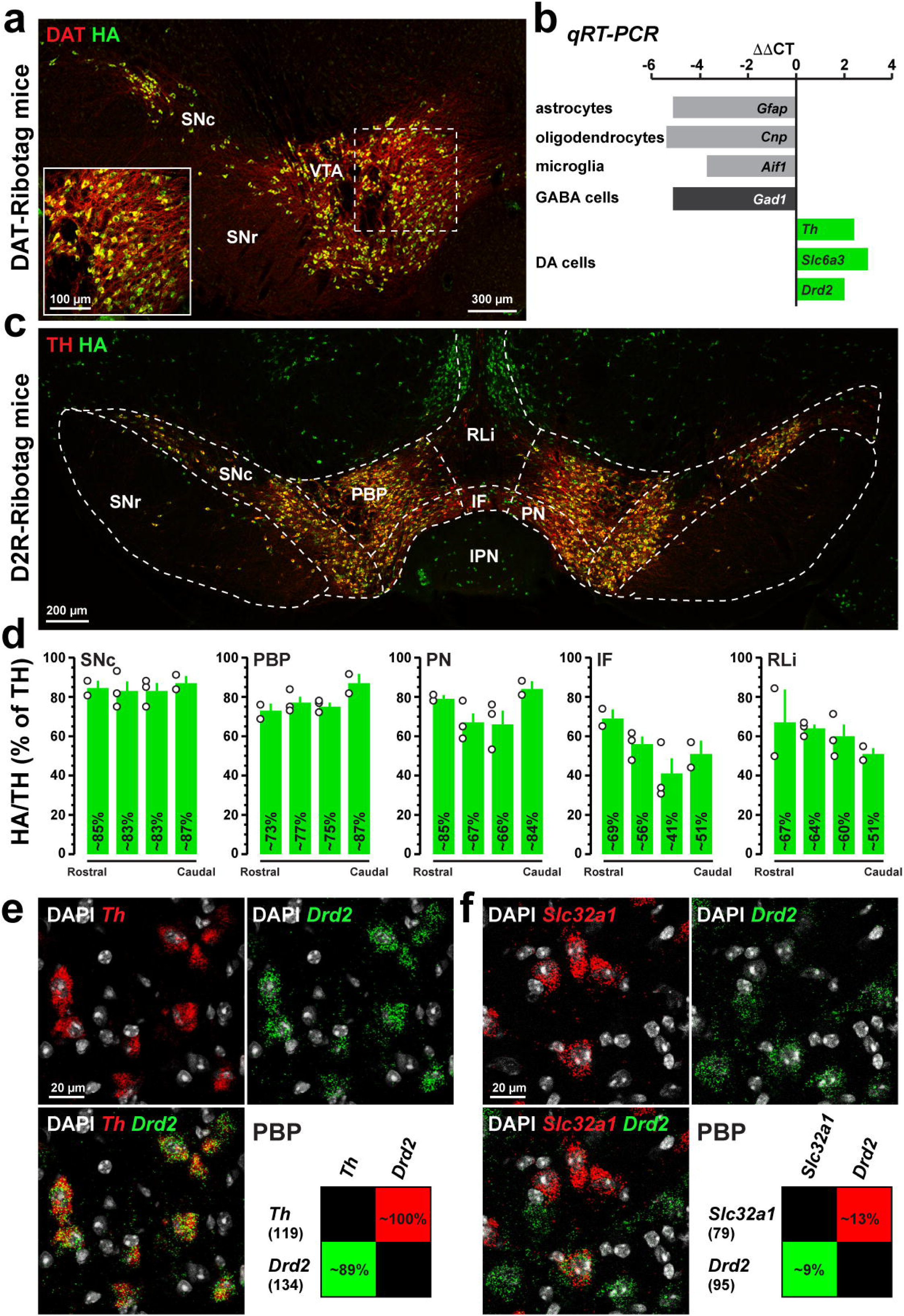
*Drd2* expression in midbrain DA-neurons. (**a**) HA (green) and TH (red) immunofluorescence in VTA DA-neurons of DAT-Ribotag mice. Note that the HA epitope fused to the ribosomal protein Rpl22 is expressed exclusively in DA-neurons identified using TH antibody (inset). (**b**) Validation by qRT-PCR (ΔΔCT) of the enrichment of VTA DA-neurons markers (*Th*, *Slc6a3*, *Drd2*) and depletion of transcripts of GABA neurons (*Gad1*) and glial cell markers including astrocytes (*Gfap*), oligodendrocytes (*Cnp*) and microglia (*Aif1*) after HA immunoprecipitation of midbrain extract compared with the input fraction (containing transcripts from all cell types). (**c**) HA (green) and TH (red) immunofluorescence in the midbrain D2R-Ribotag mice. (**d**) Analysis of HA/TH co-expression as percentage of TH-positive neurons throughout the rostro-caudal extension of SNc and VTA subareas. (**e-f**) High magnification of confocal images of coronal brain section of the VTA from C57BL/6 mouse (n = 3 mice) showing the distribution of *Drd2* (green), *Th* (red) (**e**) or *Slc32a1* (red) (**f**) expression detected with single-molecular fluorescent *in situ* hybridization. Slides were counterstained with DAPI (white). Quantification of the overlap between neurons co-expressing *Drd2*, and *Th* or *Slc32a1* in the PBP. Values in parentheses indicate the number of neurons identified for each marker (n = 3 mice). Percentages of co-labelling are represented in matrix with probes in columns among neurons labeled with probes in rows. TH: tyrosine hydroxylase; HA: hemagglutinin; SNc: substantia nigra pars compacta; SNr: substantia nigra pars reticulata; VTA: ventral tegmental area; PBP: parabrachial pigmented area; PN: paranigral nucleus; IF: interfascicular nucleus; RLi: rostral linear nucleus; IPN: interpeduncular nucleus.

#### Immunofluorescence after slices recording

After recordings, slices were fixed in 4% PFA (paraformaldehyde) during a night. Immunostaining experiments were performed as follows: free-floating VTA brain sections were incubated for 3 hours at 4°C in a blocking solution of phosphate-buffered saline (PBS) containing 3% bovine serum albumin (BSA, Sigma; A4503) (vol/vol) and 0.2% Triton X-100 (vol/vol), and then incubated during 72 hours at 4°C with a mouse anti-tyrosine hydroxylase antibody (anti-TH, Sigma, T1299), at 1:200 dilution, in PBS containing 1.5% BSA and 0.2% Triton X-100. After 3 nights, sections were rinsed with PBS, and then incubated for 6 hours at room temperature with Cy3-conjugated anti-mouse and AMCA-conjugated streptavidin (Jackson ImmunoResearch) both at 1:200 dilution in a solution of 1.5% BSA in PBS. After three rinses in PBS, slices were wet-mounted using Prolong Gold Antifade Reagent (Invitrogen, P36930). Microscopy was carried out with a fluorescent microscope, and images captured using a camera and analyzed with ImageJ. Immunoreactivity for both TH and biocytin allowed us to confirm the neurochemical phenotype of DA neurons in the VTA.

#### In situ hybridization

Brains were rapidly extracted, placed on dry ice for 5 min and stored at - 80°C. Tissue was included in an embedding medium to ensure optimal cutting temperature and then sectioned on a cryostat at 14 µm. Ventral tegmental area (VTA) coronal sections were collected onto Superfrost Plus slides (Fisher Scientific). *In situ* hybridization was carried out using the RNAscope Fluorescent Multiplex labeling kit (ACDBio catalog #323110) and the protocol previously described (Cutando et al., 2021). Probes used include Mm-Drd2-C3 (ACDBio catalog #406501-C3), Mm-Slc32a1-C2 (ACDBio catalog #319191-C2), MM-Th-C2 (ACDBio catalog #317621-C2), and Mm-Nalcn-C1 (ACDBio catalog #415165-C1). Hybridized slices were counterstained with DAPI and mounted with ProLong Diamond Antifade mounting medium (Thermo Fisher Scientific catalog #P36961). Confocal microscopy and image analyses were carried out at the Montpellier RIO imaging facility. Double-labeled images from the VTA were single confocal sections captured using sequential laser scanning confocal microscopy (Leica SP8). Values in the histograms represent co-expression as percentage of *Drd2*-expressing cells (green) and as percentage of cells expressing the other markers tested (*Th* and *Slc32a1*) (3 images in the parabrachial pigmented area per mouse, n = 3 mice).

### Polysome immunprecipitation

HA-tagged ribosome immunoprecipitation was conducted as previously described (Puighermanal et al., 2017). Briefly, the VTA from DAT-RiboTag mice (n = 5) was homogenized in 1-ml polysome buffer (50 mM Tris, pH 7.4, 100 mM KCl, 12 mM MgCl2, and 1 % NP-40, supplemented with 1 mM DTT, 1 mg/ml heparin, 100 µg/ml cycloheximide, 200 U/ml RNAseOUT, and protease inhibitors) and centrifuged at 10,000 x *g* for 10 min. Then, 100 µl of the supernatant was reserved as the input fraction. Anti-HA antibodies (5 µl/sample; Covance, #MMS-101R) were added to the remaining supernatant for overnight incubation at 4 °C, followed by incubation with protein G magnetic beads (Invitrogen, #100.04D) for another overnight period. After bead binding, samples underwent two 10-min washes with high-salt buffer (50 mM Tris, pH 7.4, 300 mM KCl, 12 mM MgCl2, 1 % NP-40, 1 mM DTT, and 100 µg/ml cycloheximide). After washing, 350 µl of Qiagen RLT buffer (supplemented with ß-Mercaptoethanol) were added to the pellets and input samples. The final RNA was extracted according to manufacturer’s instructions using a Qiagen RNeasy Micro kit and quantified on a Nanodrop 1000 spectrophotometer.

### cDNA Synthesis and Quantitative Real-Time PCR

After HA-tagged-ribosome immunoprecipitation, synthesis of cDNA and qRT-PCR were performed as previously described (Puighermanal et al., 2017). SYBR Green PCR master mix was used on the LC480 Real-Time PCR System (Roche) (n = 5 samples/group). Results are shown as linearized Cp-values, normalized to the housekeeping gene *Gapdh*, and the ΔCP method was applied to calculate fold changes. Data are expressed as the fold change between the pellet and input fractions. Primer sequences can be found in Table 1.

### Western Blotting

Western blotting was performed as previously described (Berland et al., 2022). Briefly, the brain was rapidly extracted and the striatum dissected on an ice-cold surface. Tissues were sonicated in 300 µl of lysis buffer (1% SDS-PBS containing 0.2% phosphatase inhibitors and 1% protease inhibitors) and boiled for 10 minutes. Aliquots (2.5 µl) were used for protein quantification using a BCA kit (BC Assay Protein Quantitation Kit, Interchim Uptima, Montluçon, France). Equal amounts of proteins (15 µg), supplemented with a Laemmli buffer, were loaded onto 10% polyacrylamide gels. Proteins were separated by SDS-PAGE and transferred (Trans-Blot® TurboTM, Bio-Rad) to 0.2 µm PVDF membranes (Bio-Rad, #1704156). Membranes were immunoblotted with the following antibodies: rabbit anti-VMAT2 (1:1000, Synaptic Systems, #138302), rabbit anti-TH (1:1000, Merck Millipore, #AB152), and mouse anti-β-actin (1:2000, Sigma Aldrich, #A1978). Detection was based on HRP-coupled secondary antibody binding using ECL. The secondary antibodies were: anti-mouse (1:2000, Cell Signaling Technology, #7076S) and anti-rabbit (1:2000, Cell Signaling Technology, #7074S). Membranes were imaged using the Amersham Imager 680. Quantifications were performed using the ImageJ software.

### Microdialysis and Dopamine assay

Surgery and perfusion procedures were performed as previously described (Cathala et al., 2022) with minor modifications. For all experiments, the microdialysis probe (CMA/11, cuprophan, 240µm outer diameter, Carnegie Medicin, Phymep, France) was 1 mm length for the NAc. Mice was anesthetized with 3% isoflurane, and placed in a stereotaxic frame. The microdialysis probe was implanted according to the atlas of Paxinos and Franklin (2001) into the nucleus accumbens (coordinates, in mm, relative to the Bregma point, AP = 1.35, L = -0.5, V = -5.2). After the surgery, the percentage of isoflurane was adjusted to 1.5% until the end of the experiment. Probes were perfused at a constant flow rate (1 μl/min), by means of a microperfusion pump (CMA 111, Carnegie Medicin, Phymep), with aCSF containing (in mM): 147 NaCl, 2.7 KCl, 1.2 CaCl2, 1 MgCl2 in phosphate buffer, pH 7.4. Dialysates were collected in a refrigerated fraction collector (MAB 85 Microbiotech, Phymep) 120 min after the beginning of the perfusion (stabilization period) and every 15 min. DA outflow was monitored during 60 min. After collection, dialysate samples were immediately analyzed with a high-performance liquid chromatography apparatus (Alexys UHPLC/ECD Neurotransmitter Analyzer, Antec, The Netherlands), equipped with an autosampler (AS 110 UHPLC cool 6-PV, Antec), as previously described (Cathala et al., 2022). The mobile phase [containing (in mM) 100 phosphoric acid, 100 citric acid, 0.1 EDTA.2H2O, 4.6 octanesulfonic acid.NaCl plus 5% acetonitrile, adjusted to pH 6.0 with NaOH solution (50%)] was delivered at 0.070 ml/min flow rate with a LC 110S pump (Antec) through an Acquity UPLC BEH column (C18; 1 x 100 mm, particle size 1.7 μm; Waters, Saint-Quentin en Yvelynes, France). Detection of DA was carried out with an electrochemical detector (DECADE II, Antec) with a VT-03 glassy carbon electrode (Antec) set at +460 mV versus Ag/AgCl. Output signals were recorded on a computer (Clarity, Antec). Under these conditions, the retention time for DA was 3-3.5 min. and the sensitivity was 50 pM with a signal/noise ratio of 3:1. At the end of each experiment, the animal was deeply anesthetized with a pentobarbital overdose (Exagon, 200mg/kg, Centravet), and its brain was removed and fixed in NaCl (0.9%)/paraformaldehyde solution (10%). Probe location into the NAc was determined histologically on serial coronal sections (60 µm), and only data obtained from animals with correctly implanted probes were included in the results.

### Electrophysiology

#### In vivo electrophysiology on anesthetized animals–juxtacellular recordings

Adult male mice (8-12 weeks) were deeply anesthetized with chloral hydrate (8%) (400 mg/kg, i.p.), supplemented as required to maintain optimal anesthesia throughout the experiment. Mice were then placed in a stereotaxic frame (David Kopf), the scalp was opened and a cranial window was drilled in the skull above the location of the VTA (coordinates: 3.1 ± 3 mm posterior to bregma, 0.4 to 0.5 mm lateral to the midline, 3.9 to 5 mm ventral from the brain). Recording electrodes were pulled from borosilicate glass capillaries (Harvard Apparatus, with outer and inner diameters of 1.50 and 1.17 mm, respectively) with a Narishige electrode puller. The tips were broken under microscope control and filled with 0.5% sodium acetate. Electrodes had tip diameters of 1-2 μm and impedances of 6–9 MΩ. A reference electrode was placed in the subcutaneous tissue. The recording electrodes were lowered vertically through the hole with a micro drive. Electrical signals were amplified by a high-impedance amplifier (Axon Instruments) and monitored through an audio monitor (A.M. Systems Inc.). The unit activity was digitized at 12.5 kHz and recorded using Spike2 software (Cambridge Electronic Design). Individual electrode tracks were separated from one another by at least 0.1 mm in the medio-lateral axis. The electrophysiological characteristics of DA-neurons were analyzed in the active cells encountered when passing the microelectrode in a stereotaxically defined block of brain tissue corresponding to the coordinates of the VTA. Extracellular identification of putative DA-neurons was based on their location as well as on the set of unique electrophysiological properties that distinguish dopamine from non-dopamine neurons *in vivo*: (i) a typical triphasic action potential (AP) with a marked negative deflection; (ii) a long duration (>2.0 ms); (iii) an action potential width from start to negative trough >1.1 ms; (iv) a slow firing rate (<10 Hz and >1 Hz). Electrophysiological recordings were analyzed using the R software. DA cell firing was analyzed with respect to the average firing rate and the percentage of spikes within bursts (% SWB, number of spikes within burst divided by total number of spikes). Bursts are classically identified as discrete events consisting of a sequence of spikes with (i) a burst onset defined by two consecutive spikes within an interval <80 ms and (ii) the end of a burst defined by an inter-spike interval >160 ms. To study the response of DA-neurons to nociceptive stimuli, three tail pinches (duration 5 s, interval 1 min) were applied manually. To record the response of silent, putative DA-neurons to nociceptive stimuli, microelectrodes were slowly lowered (4 µm steps) into the VTA until an increase in resistance indicated proximity to a cell. Next, three manual tail pinches (5-second duration, 1-minute intervals) were applied to induce AP. The percentage of mean firing frequency variation induced by pinch was calculated by comparing AP frequency during the 5-second pinch period to the baseline frequency measured 10 seconds prior. For each neuron, the average percentage change in mean firing frequency was obtained from the three pinch trials. Neurons with a positive change in mean firing frequency were classified as excited (Type 1) in response to the pinch, while those with a negative change were classified as inhibited (Type 2). To estimate the maximum firing variation caused by the pinch, we calculated the instantaneous firing frequency in 3-second sliding windows. The maximal variation of instantaneous frequency from the baseline (measured 10 seconds before the pinch) was calculated during the 5-second pinch period. For each neuron, the maximum firing variation was obtained by averaging the peak variation across three separate pinches. Both the mean and maximum firing frequency variation induced by pinch are expressed as percentage changes from baseline (mean ± SEM). Experimenters were blinded to the mice’s genotype during recording.

#### Ex vivo electrophysiology-patch clamp recordings

Mice were deeply anesthetized by an intraperitoneal injection of a mix of ketamine (150 mg/kg Virbac 1000) and xylazine (60 mg/kg, Rompun 2%, Elanco). Coronal midbrain sections (250 μm) were sliced with a Compresstome (VF-200, Precisionary Instruments) after intracardial perfusion of cold (4°C) sucrose-based artificial cerebrospinal fluid containing (in mM): 125 NaCl, 2.5 KCl, 1.25 NaH_2_PO_4_, 26 NaHCO_3,_ 5,9 MgCl_2_, 25 sucrose, 2.5 glucose, 1 kynurenate (pH 7.2, 325 mOsm). After 8 minutes at 37°C for recovery, slices were transferred into oxygenated artificial cerebrospinal fluid (ACSF) containing (in mM): 125 NaCl, 2.5 KCl, 1.25 NaH_2_PO_4_, 26 NaHCO_3,_ 2 CaCl_2_, 1 MgCl_2_, 15 sucrose, 10 glucose (pH 7.2, 325 mOsm) at room temperature for the rest of the day. Slices were individually transferred to a recording chamber continuously perfused at 2 mL/minute with oxygenated ACSF. Patch pipettes (4-6 MΩ) were pulled from thin wall borosilicate glass (G150TF-3, Warner Instruments) with a micropipette puller (P-87, Sutter Instruments Co.) and filled with a potassium gluconate-based intracellular solution containing (in mM): 135 K-gluconate, 10 HEPES, 0.1 EGTA, 5 KCl, 2 MgCl_2_, 2 ATP-Mg, 0.2 GTP-Na, and biocytin 2 mg/mL (pH adjusted to 7.2). Neurons were visualized using an upright microscope coupled with a dodt gradient contrast imaging, and illuminated with a white light source (Scientifica). Whole-cell recordings were performed with a patch-clamp amplifier (Axoclamp 200B, Molecular Devices) connected to a Digidata (1550 LowNoise acquisition system, Molecular Devices). Signals were low-pass filtered (Bessel, 2 kHz) and collected at 10 kHz using the data acquisition software pClamp 10.5 (Molecular Devices). VTA location was identified under microscope. Identification of DA-neurons was performed by location and by their electrophysiological properties (width and shape of action potential (AP) and after hyperpolarization (AHP)). Recorded neurons in *patch-clamp* experiments were filled with biocytin in order to validate presence of Tyrosine hydroxylase (TH) by immunofluorescence as an indicator of DA-neurons.

### Auditory brainstem response recordings and distortion product otoacoustic emission recordings

Auditory brainstem responses (ABR) were carried out under anesthesia with Rompun 2% (3 mg/kg) and Zoletil 50 (40 mg/kg) in a Faraday shielded anechoic soundproof. Rectal temperature was measured with a thermistor probe and maintained at 38.5°C ± 1 using a heater under-blanket (Homeothermic Blanket Systems, Harvard Apparatus). The acoustical stimuli consisted of 9-ms tone bursts, with a plateau and a 1-ms rise/fall time, a delivery rate from 11/sec to 20/sec of alternate polarity by a JBL 2426H loudspeaker in a calibrated free field. Stimuli were generated and data acquired using Matlab (MathWorks) and LabView (National Instruments) software. The difference potential between vertex and mastoid subcutaneous needles was amplified (2500 times, VIP-20 amplifier), sampled (at a rate of 50 kHz), filtered (bandwidth of 0.3-3 kHz), and averaged (500 to 700 times). Data were displayed using LabView software and stored on a computer (Dell T7400). ABR thresholds were defined as the lowest sound intensity that elicits a clearly distinguishable response. For distortion product otoacoustic emission (DPOAE) recordings, an ER-10C S/N 2528 probe (Etymotic Research), consisting of two emitters and one microphone, was inserted in the left external auditory canal. Stimuli were two equilevel (65 dB SPL) primary tones of frequency f_1_ and f_2_ with a constant f_2_/f_1_ ratio of 1.2. The distortion 2f_1_-f_2_ was extracted from the ear canal sound pressure and processed by the HearID auditory diagnostic system (Mimosa Acoustic) on a computer (Hewlett Packard). The probe was self-calibrated for the two stimulating tones before each recording. f_1_ and f_2_ were presented simultaneously, sweeping f_2_ from 20 to 2 kHz in quarter octave steps. For each frequency, the distortion product 2f_1_-f_2_ and the neighbouring noise amplitude levels were measured and expressed as a function of f_2_.

### Behaviors

#### Locomotor activity

Horizontal and vertical activity was measured for 60 min in a non-stressful environment in a circular corridor (Imetronic, Pessac, France) for 60 min (Puighermanal et al., 2020).

#### Elevated plus maze

Elevated plus maze was performed as previously described (Puighermanal et al., 2020). The number of entries and time spent in the closed arms (4 paws within closed arm) or open arms (4 paws within open arms) were calculated during 5 min.

#### Hotplate test

Thermal nociception was assessed using a hotplate. Mice were place in plexiglass chamber (17 x 17 x 25) on a hotplate at 52°C. The latency to lick front paws was recorded using a stopwatch.

#### Discriminative Auditory threat conditioning

The experiment was carried out in soundproof fear conditioning apparatus (Imetronic, Pessac, France) as previously described (Castell et al., 2024). Discriminative the auditory threat conditioning was carried out as follows: Day 1, *habituation*: after 5 min of exploration in box A or B, two different tones (CS+ and CS-; 30 sec 4 times each) were randomly presented to the mice: *conditioning*: the same day, mice were placed on the same box as for the habituation and after 5 min of exploration they received 5 pairings of the CS+ (30 sec) with the foot-shock at the last second of the CS+ (1 sec, 0.6 mA), and 5 presentations of the CS-alone (30 sec). Interval between each tone was a random time between 20 and 180 sec. Day 2-4: animals underwent the test (day 2) and two extinction (day3-4) sessions on a different box during 3 consecutive days. On these sessions, animals received 12 presentations of the CS+ (30 sec) and 4 presentations of the CS-. The intertrial interval in all sessions was between 20 and 180 sec. In the habituation and conditioning, cages were cleaned with 70% ethanol and for the test and extinction sessions with 1% acetic acid. The meaning of each tone (pairing or not with the foot-shock) was counterbalanced between animals and genotypes. Freezing behavior was recorded using a tight infrared frame. The threshold for considering freezing behavior was set up at 2 sec. The first 10 min of habituation were used to assess the basal freezing. Data are presented as the probability to evoke freezing behavior during CS+ and CS-presentation.

#### Discriminative Active avoidance

The experiment was carried out in a soundproof shuttle box (Imetronic, Pessac, France) as previously described (Gangarossa et al., 2019). The apparatus is made of two equal compartments (20 x 20 cm) separated by an opened door. Both compartments have a metal grid on the floor, independent houselights and one infrared beam frame on each compartment. Day 1: mice were subjected to a habituation session consisting of 10 presentations of two different tones, a CS- and CS+ lasting 13 sec maximum. Subjects can stop both CSs by shuttling from one compartment to another. Day 2-4: mice were subjected to the training sessions once per day during which they were exposed to CS+/CS-in a pseudo-random manner. Again, the CS (+ or -) stops whenever the mice cross from one compartment to another. CS-last 8 sec maximum. CS+ last 13 sec maximum, but after 8 sec the unconditioned stimulus (0.4 or 0.7 mA foot-shock coinciding with the last 5 sec of the CS+presentation) is delivered until the mice shuttles or for a maximum of 5 sec. During the inter-trial, the animal was free to cross between compartments. The probability of avoidances was used as an index of learning.

### Statistical analysis

All statistical analyses were done using the R software with home-made routines. Results were plotted as mean ± SEM. The total number (n) of observations in each group and the statistical tests used for comparisons between groups or within groups are indicated directly on the figures or Supplemental material. Comparisons between distributions were performed with Kolmogorov-Smirnov test. Comparisons between means were performed with parametric tests such as Student’s t-test, or two-way ANOVA for comparing two groups when parameters followed a normal distribution (Shapiro-Wilk normality test with p > 0.05), or Wilcoxon non-parametric test if the distribution was skewed. ABR were analyzed with Mann–Whitney Wilcoxon’s test. Western Blots and microdialysis experiments were analyzed with unpaired Student t-test. Behaviors were analyzed with two-way repeated measure ANOVA, unpaired or paired Student t-test as detailed in Supplemental Table 2. Statistical significance was set at p < 0.05 (*), p < 0.01 (**), or p < 0.001 (***), or p > 0.05 was considered not to be statistically significant.

## Results

### Distribution of *Drd2* among midbrain DA-neurons

We analyzed the expression of *Drd2* transcripts from tagged ribosome-bound mRNAs of VTA extracts of DAT-Ribotag mice. First, we confirmed that hemagglutinin (HA) expression, which labels the tagged ribosomal protein Rpl22, was restricted to DAT-positive neurons using double immunofluorescence (**Figure 1a**). Taking advantage of the Ribotag approach, we observed that, in contrast to gene expression markers of astrocytes (*Gfap*), oligodendrocytes (*Cnp*), microglia (*Aif1*), and GABAergic cells (*Gad1*), markers o DA-neurons (*Th, Slc6a3* and *Drd2*) were significantly enriched following immunoprecipitation of tagged ribosomes from VTA homogenates (**Figure 1b**). We then assessed the co-expression of TH and D2R throughout the rostro-caudal extent of the SNc and VTA subregions, including the parabrachial pigmented area (PBP), paranigral nucleus (PN), interfascicular nucleus (IF), and rostral linear nucleus (RLi), using D2R-Ribotag male mice (**Figure 1c-d**). Our analysis revealed that the percentage of TH/HA co-immunoreactive neurons was high in the SNc (∼84%), PBP (∼77%), and PN (∼71%), with slightly lower percentages in the midline VTA subareas [IF (∼51%) and RLi (∼58%)] (**Figure 1c-d**). Finally, the enrichment of *Drd2* transcripts in VTA DA-neurons (exemplified in the PBP) was confirmed through single-molecule fluorescent *in situ* hybridization. This analysis revealed that *Drd2* was present in a high proportion of *Th*-expressing cells (∼89%) (**Figure 1e**). Interestingly, *Drd2* transcripts were also detected in ∼9% of cells expressing the gene encoding the vesicular GABA transporter (*Slc32a1*), indicating that D2R is also present in a subset of VTA GABA-neurons (**Figure 1f**).

### Enhanced activity of VTA DA-neurons in response to salient aversive stimuli in mice lacking autoD2R

To investigate the role of autoD2R, we generated mice lacking *Drd2* specifically in midbrain DA-neurons by crossing *Drd2^loxP/loxP^* mice with the *Slc6a3-Cre* mouse line (referred to as *Drd2^Slc6a3^* mice) (**Supplemental Figure 1**). As expected, D2R immunoreactivity was reduced in the striatum of *Drd2^Slc6a3^* mice as compared to control mice (**Supplemental Figure 1a**), whereas no alterations in the expression levels of TH and VMAT2 were detected (**Supplemental Figure 1b-d**), nor reduced extracellular levels of basal DA were measured in the Acb (**Supplemental Figure 1e)**. We next assessed the consequences of this selective deletion on the firing patterns of midbrain DA-neurons using *in vivo* juxtacellular recordings in anesthetized male mice (**Figure 2a**). Analyses of firing frequency and the percentage of spikes within burst (%SWB) of VTA DA-neurons revealed no differences between control and *Drd2^Slc6a3^* male mice, suggesting that the absence of autoD2R in DA-neurons does not affect their firing patterns (**Figure 2a**).

**Figure 2:**
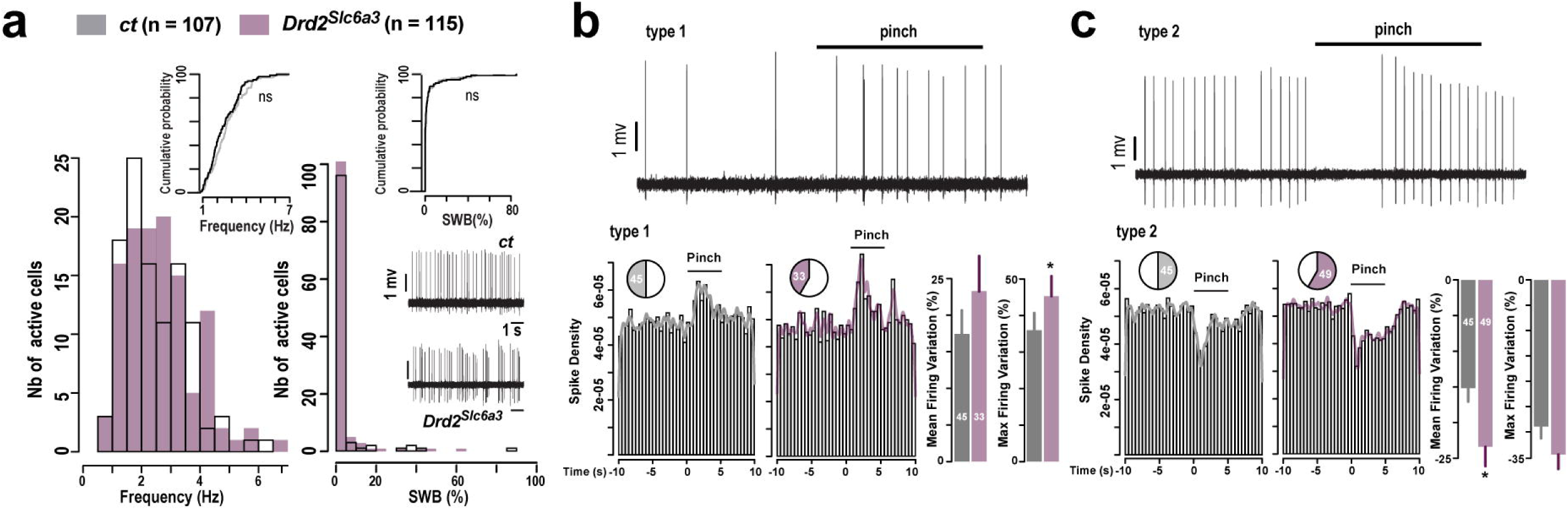
*In vivo* VTA DA-neurons activity in response to salient aversive stimuli in mice lacking *Drd2* in DA-neurons. (**a**) Distribution of firing frequency (left) and percentage of SWB (Rigth) of VTA DA neurons spontaneous activity in *ct* (grey) and *Drd2^Slc6a3^* (purple) (n = 107 *ct*, n = 115 *Drd2^Slc6a3^*). (**b**) Pinch-activated VTA DA neurons in *ct* (grey) and *Drd2^Slc6a3^* (purple). Left: Cumulative spikes density 10 sec before and after the pinch for all the tail pinch (3 pinch per neurons, n = 45 neurons of 10 mices in *ct*, n = 33 neurons of 10 mices in *Drd2^Slc6a3^*). Right: Percentage of mean firing frequency variation and maximum firing frequency variation of DA neurons induced by pinch (n = 45 neurons of 11 mice in *ct*, n = 33 neurons of 11 mice in *Drd2^Slc6a3^*). (**c**) Same as (**b**) for pinch-inhibited VTA DA neurons (n = 45 neurons of 11 mice in *ct*, n = 49 neurons of 11 mice in *Drd2^Slc6a3^*). Statistical analysis detailed in Supplemental Table (2a-c)

Given that VTA DA-neurons show either inhibition or a phasic increases in firing rate in response to aversive stimuli (Brischoux et al., 2009; Ungless et al., 2004), we aimed at determining whether these responses were altered in *Drd2^Slc6a3^* mice. We measured the firing rate of VTA DA-neurons following repeated tail pinches, used here as salient aversive stimuli. In control male mice, VTA DA-neurons were classified into two categories: those exhibiting excitation (type 1) and those displaying rapid inhibition (type 2) (**Figure 2b, c**). Interestingly, both excitatory and inhibitory responses were significantly enhanced in *Drd2^Slc6a3^* male mice as compared to controls, suggesting that autoD2Rs scale VTA DA-neurons firing rate in response to aversive stimuli.

### Tonic activity of VTA DA-neurons is impaired in mice lacking *Nalcn* in DA-neurons

Recent evidence suggest that suggests that D2Rs exert a key mechanistic control over midbrain DA-neurons excitability through a G_i/o_-dependent inhibition of the non-selective sodium leak channel NALCN (Cobb-Lewis et al., 2023; Philippart & Khaliq, 2018; Um et al., 2021). Using *in situ* hybridization, we first assessed the percentage of cells expressing *Nalcn* transcripts in VTA autoDA-neurons and found that *Nalcn* transcripts were detected in ∼100% of autoDA-neurons (**Figure 3a-c**). Of note, *Nalcn* transcripts were also found in VTA GABA-neurons, thus illustrating the ubiquitous expression of NALCN in VTA (**Figure 3b-c**).

**Figure 3:**
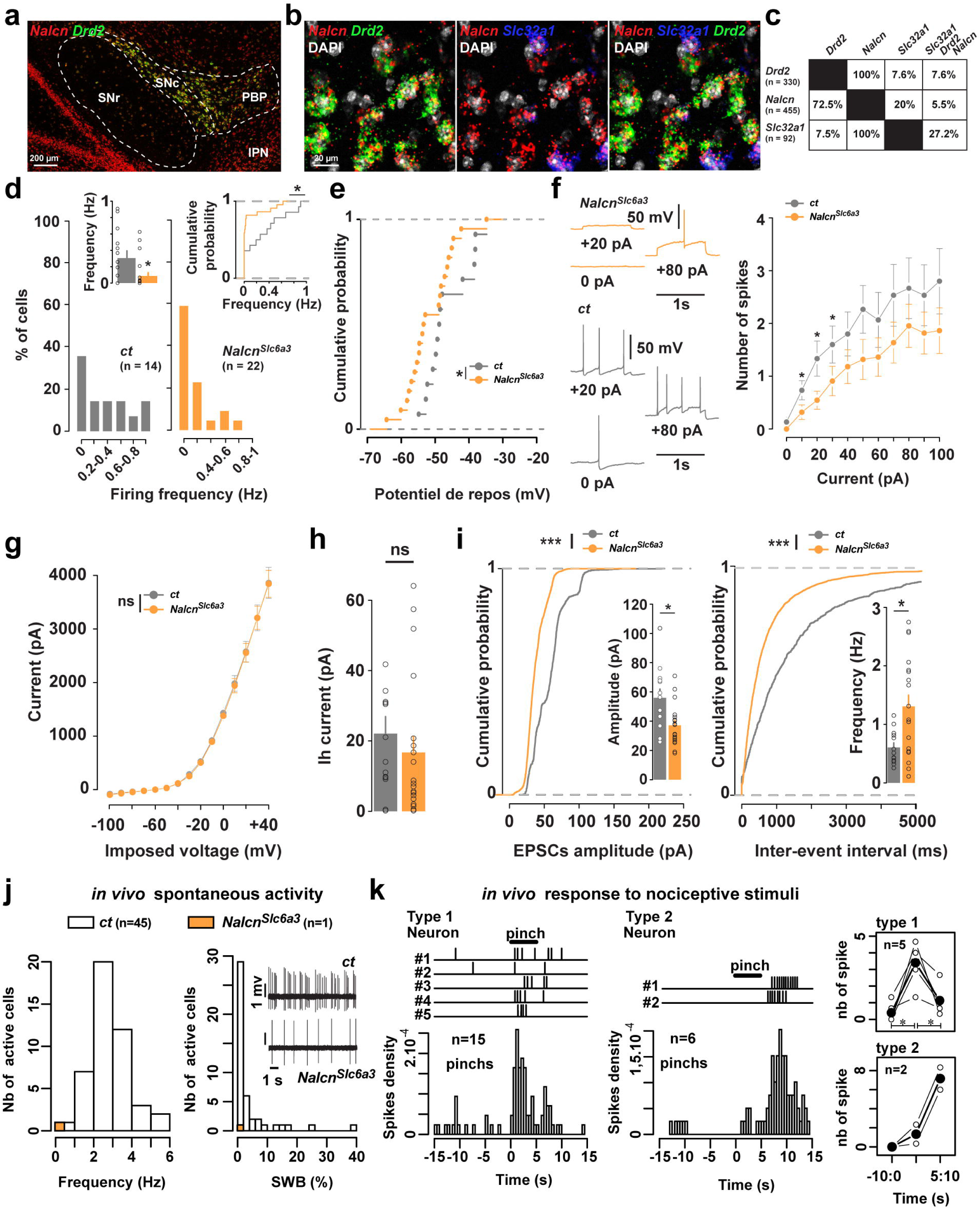
*In vitro* and *in vivo* characterization of electrophysiological properties of VTA DA-neurons lacking *Nalcn*. (**a**) Single fluorescence *in situ* hybridization for *Nalcn* and *Drd2* transcripts in the ventral midbrain. (**b**) High magnification of confocal images of coronal brain section of the VTA from C57BL/6 mouse (n = 3 mice) showing *Drd2* (green), *Nalcn* (red) and *Slc32a1* (blue) mRNA expression detected with single-molecular fluorescent *in situ* hybridization. Slides were counterstained with DAPI (white). (**c**) Quantification of the overlap between neurons co-expressing *Drd2*, *Nalcn* and *Slc32a1* in the PBP. Values in parentheses indicate the number of neurons identified for each marker (n = 3 mice). Percentages of co-labelling are represented in matrix with probes in columns among neurons labeled with probes in rows. (**d**) Histogram density of firing rates of *ct* (grey; n = 14 neurons; n = 2 mice) and *Nalcn^Slc6a3^* (orange; n = 22 neurons; n = 3 mice) mice, with corresponding cumulative probability (inset). (**e**) Cumulative probability of resting membrane potential of *ct* (grey; n = 14 neurons; n = 2 mice) and *Nalcn^Slc6a3^* (orange; n = 22 neurons; n = 3 mice) mice. (**f**) Example traces of firing of VTA DA-neurons of *ct* and *Nalcn^Slc6a3^* mice after current injections (0, +20, +80 pA). FI-curve (frequency-current relationship) of *ct* (grey; n = 13 neurons; n = 2 mice) and *Nalcn^Slc6a3^* (orange; n = 22 neurons; n = 3 mice) mice. Lower excitability of VTA DA neurons of *Nalcn^Slc6a3^* mice for no or small currents (0, 10, 20, 30 pA) compared to DA neurons of *ct* mice. (**g**) IV-curve (current-voltage relationship) of *ct* (grey; n = 14 neurons; n = 2 mice) and *Nalcn^Slc6a3^* (orange; n = 22 neurons; n = 3 mice) mice. (**h**) Ih-current induced by hyperpolarizing steps in VTA DA neurons of *ct* (grey; n = 14 neurons; n = 2 mice) and *Nalcn^Slc6a3^* (orange; n = 20 neurons; n = 3 mice) mice. (**i**) Cumulative probability of spontaneous post-synaptic excitatory currents (sEPSC) amplitude (left) and frequency (right) recorded from *ct* (grey; n = 13 neurons; n = 2 mice) and *Nalcn^Slc6a3^* (orange; n = 19 neurons; n = 3 mice) mice. Insets: bar graphs of the means obtained from sEPSC amplitude (left) or frequency (right;data represent means ± SEM). (**j**) *In vivo* juxtacellular recordings of putative DA neurons on anesthetized animals. Distribution of spontaneous firing frequency among active putative DA neurons in the VTA of *ct* (grey, n = 45 neurons of 2 mice) and *Nalcn^Slc6a3^* (orange, n = 1 neuron of 2 mice) mice. pDA of *Nalcn^Slc6a3^* mice are silent except one neuron, compared to putative DA-neurons of *ct* mice showing frequency up to 6 Hz and bursting activity. (**k**) *In vivo* response to nociceptive stimuli of silent putative DA neurons on anesthetized *Nalcn^Slc6a3^* mice. Left: raster plot of AP evoked during a 5 s-duration tail pinch in 5 neurons (type 1) and cumulative density of AP 15s before and after tail pinch (15 pinch, 3 pinch per neuron). Middle: raster plot of AP evoked at the end a 5 s-duration tail pinch in 2 neurons (type 2) and cumulative density of AP 15 s before and after tail pinch (6 pinch, 3 pinch per neuron). Right: mean number of AP, 10 s before, during or 5 s after the pinch on Type 1 (upper) or Type 2 (lower) neurons. White dot: mean number of AP per neuron for 3 pinch. Black dot: mean number of AP on 5 type 1 and 2 type 2 neurons. *p < 0.05, ***p < 0.001. Statistical analysis detailed in Supplemental Table (3d-i).

To explore the role of NALCN in regulating the firing patterns of midbrain DA-neurons, we generated mice with a selective deletion of *Nalcn* in midbrain DA-neurons by crossing *Nalcn^loxP/loxP^* mice (Flourakis et al., 2015) with the *Slc6a3-Cre* mouse line (referred to as *Nalcn^Slc6a3^* mice). Although no alterations in D2R, TH and VMAT2 levels were found in the striatum of *Nalcn^Slc6a3^* mice as compared to controls (**Supplemental Figure 2a-d**), we observed a significant decrease of basal DA levels in the Acb of *Nalcn^Slc6a3^* mice, suggesting that activity of VTA DA-neurons might be impaired (**Supplemental Figure 2e**). To probe this hypothesis, we compared the pacemaker activity of these neurons between control and *Nalcn^Slc6a3^* male mice using *ex vivo* patch-clamp recordings. Consistent with previous observations (Philippart & Khaliq, 2018), VTA DA-neurons from *Nalcn^Slc6a3^* male mice exhibited reduced spontaneous activity (mean *ct* = 0.3 Hz, mean *Nalcn^Slc6a3^* = 0.08 Hz, **Figure 3d**) and a more hyperpolarized resting membrane potential as compared to controls (mean *ct* = - 46 mV, mean *Nalcn^Slc6a3^* = - 51 mV, **Figure 3e**). Regarding the membrane properties, the current-response curve indicated decreased excitability of DA-neurons for small currents in *Nalcn^Slc6a3^* compared to control male mice (**Figure 3f**), although no differences were observed in the I-V relationship (**Figure 3g**) or Ih currents between the two genotypes (mean *ct* = 22 pA, mean *Nalcn^Slc6a3^* =17 pA, **Figure 3h**). We then investigated the glutamatergic drive of DA-neurons and found that spontaneous EPSCs had reduced amplitude but higher frequency in *Nalcn^Slc6a3^* mice compared to controls (for amplitudes: mean ct = 56 pA, mean *Nalcn^Slc6a3^* = 37 pA, **Figure 3i**). These results suggest that *Nalcn* deletion may also reduce the *in vivo* activity of VTA DA-neurons.

To test this, we recorded *in vivo* the activity of VTA DA-neurons using juxtacellular recordings in anesthetized male mice. Strikingly, while we recorded a substantial number of active cells in control mice (n = 45, on 2 mice), no spontaneously active neurons (except one, below 1Hz, on 3 mice) were found in *Nalcn^Slc6a3^* male mice, suggesting that, in anesthetized mice, the majority of DA-neurons are silent (**Figure 3j**). However, following repeated tail pinches, we were able to evoke excitatory responses in seven previously silent putative DA-neurons across 2 *Nalcn^Slc6a3^* male mice. Five neurons responded by firing action potentials (APs) during the stimulus (type1), while two fired APs at the end of the stimulus (Type 2) (**Figure 3k**). Together, these results indicate that the response of VTA DA-neurons to aversive stimuli is independently of D2R/G_i/o_-mediated inhibition of NALCN.

### Enhanced reactivity to threat-predicting cues in mice lacking *Drd2* in DA-neurons

Next, we investigated whether D2R/NALCN signaling in midbrain DA-neurons functionally modulates the motivational salience of discrete auditory cues predicting a threat. Control, *Drd2^Slc6a3^* and *Nalcn^Slc6a3^* male and female mice were subjected to discriminative auditory threat conditioning (**Figure 4a and Supplemental Figure 3**). On the conditioning day (day 1) both lines of control mice and *Nalcn^Slc6a3^* mice expressed similar levels of freezing in response to both CS+ and CS-(**Figure 4b-d, upper panels and Supplemental Figure 3a-c, upper panels**). In contrast, *Drd2^Slc6a3^* male mice, but not females, displayed significantly higher freezing responses to the CS+ compared to CS-, suggesting that autoD2R plays a key role in the discrimination between CS+ and CS-(**Figure 4b, upper panel and Supplemental Figure 3a, upper panel**). Of note, the observed differences in males were not attributed to changes in spontaneous locomotion and exploratory drive (**Supplemental Figure 4**), approach-avoidance behaviors (**Supplemental Figure 5**), or altered sensory systems (hearing and nociception), since *Drd2^Slc6a3^* male mice showed similar auditory thresholds (**Supplemental Figure 6**) and nociceptive thresholds (**Supplemental Figure 7**) as compared to controls. On the test day (day 2), control and *Drd2^Slc6a3^* male mice, but not females, displayed higher freezing responses to the CS+ compared to the CS-(**Figure 4b, middle panels** and **Supplemental Figure 3a, middle panels**). There were no differences in freezing responses between control and *Drd2^Slc6a3^* mice indicating that the enhanced discrimination observed in *Drd2^Slc6a3^* mice did not translate into higher overall freezing (**Figure 4b, middle panels**). In contrast, opposite sex-biased regulation was observed in *Nalcn2^Slc6a3^* mice. Indeed, control and *Nalcn2^Slc6a3^* female mice, but not males, equally discriminated between auditory cues predicting (CS+) or not (CS-) threat (**Figure 4c, middle panels** and **Supplemental Figure 3b, middle panels**). On the other hand, at higher-intensity shock (0.7 mA), discrimination was only significant in both male and female control mice **Figure 4d, middle panels** and **Supplemental Figure 3c, middle panels**). Following repeated CS+ presentation, all groups of male mice – including control, *Drd2^Slc6a3^* and *Nalcn^Slc6a3^* mice – exhibted a gradual decrease in freezing responses, indicating that extinction was not impaired (**Figure 4b-d, lower panels**). However, only *Drd2^Slc6a3^* male mice maintain a higher level of discrimination between the CS+ and CS-as compared to control male mice, suggesting that enhanced discrimination may slow down the extinction process (**Figure 4b, lower panels**).

**Figure 4:**
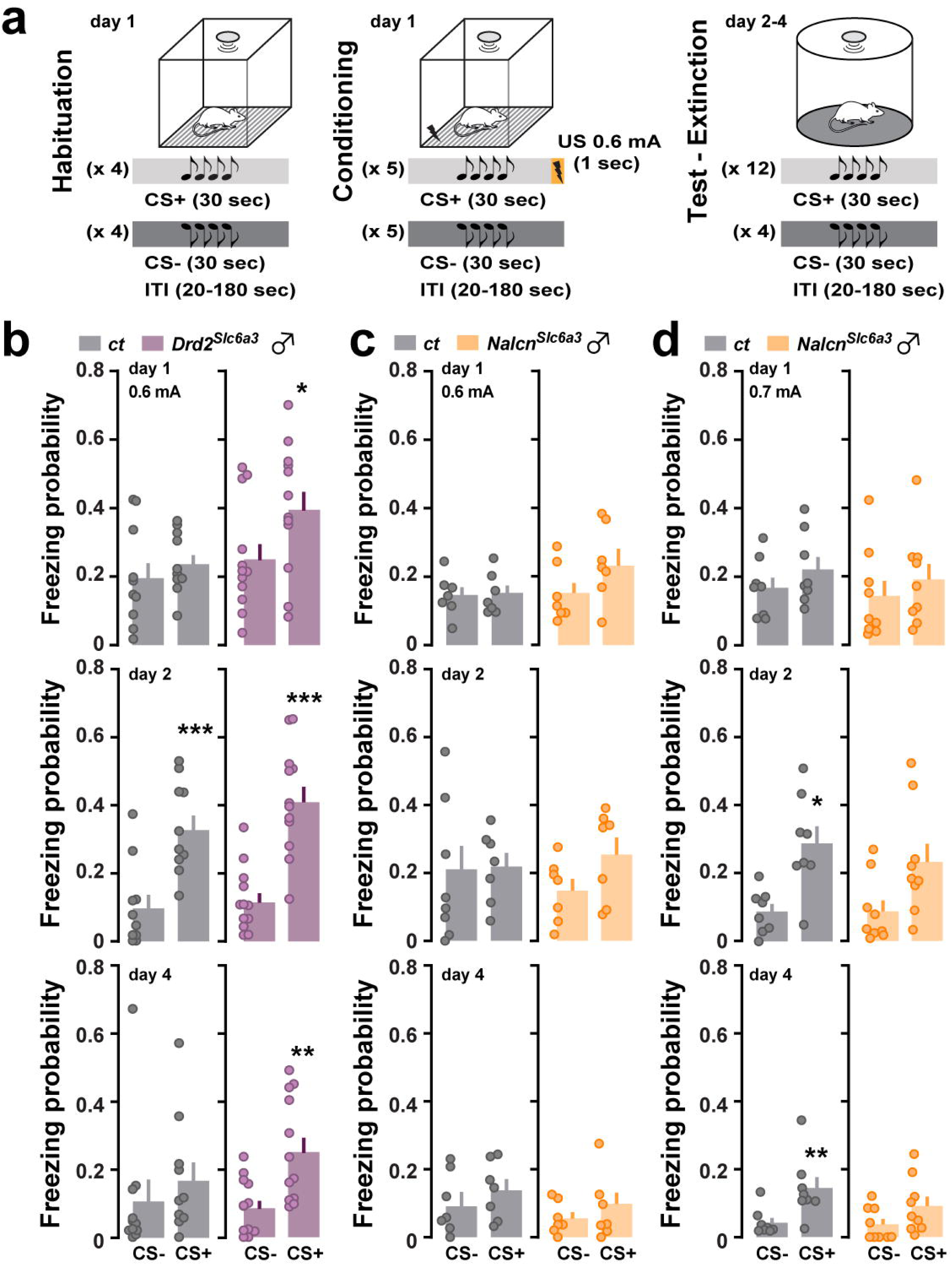
Discriminative learning to threat-predicting cues in male mice lacking *Drd2* or *Nalcn* in DA-neurons. (**a**) Schematic cartoon describing the protocol used to evaluate discriminative auditory threat conditioning. (**b**) Probability of CS- and CS+ to evoke freezing responses in *ct* (grey) and *Drd2^Slc6a3^* (purple) mice during the conditioning (day 1, upper panels), the test (day 2, middle panels) and after extinction (day4, lower panels) (n = 10 *ct*, n = 12 *Drd2^Slc6a3^*). (**c**) Similar to (**b**) in *ct* (grey) and *Nalcn^Slc6a3^* (orange) mice (0.6 mA: n = 7 *ct*, n = 7 *Nalcn^Slc6a3^*; 0.7 mA: n = 8 *ct*, n = 9 *Nalcn^Slc6a3^*). Results are analyzed using paired *t*-test. CS-vs CS+, *p < 0.05, **p < 0.01. Statistical analysis detailed in Supplemental Table (4b-d).

We then explored whether autoD2Rs in midbrain DA-neurons also regulates the reactivity to threat-predicting cues during dynamic defensive behaviors, such as avoidance. Control, *Drd2^Slc6a3^* and *Nalcn^Slc6a3^* mice were subjected to discriminative active avoidance learning (**Figure 5a**). Male and female mice from both genotypes were trained using low-intensity shocks (0.4 mA) over 3 consecutive days (**Figure 5, Supplemental Figure 8**). As expected, during the first session, *Drd2^Slc6a3^* and *Nalcn^Slc6a3^* male mice exhibited escape behaviors, characterized by low avoidance probability, regardless of the shock intensity (**Figure 5b-c and Supplemental Figure 8**). Across subsequent sessions, control, *Drd2^Slc6a3^* and *Nalcn^Slc6a3^* male mice gradually learned to avoid the threat following CS+ presentation and discriminated between CS+ and CS-(**Figure 5b-c and Supplemental Figure S8**). However, at the low (0.4 mA), but not high-intensity shock (0.7 mA), discriminative learning was enhanced in *Drd2^Slc6a3^* male mice, but not in *Nalcn^Slc6a3^* mice, as evidenced by higher avoidance probability and discriminative performance compared to control mice (**Figure 5b-e and Supplemental Figure 8**). Together, these findings suggest that autoD2Rs in midbrain DA-neurons scale the reactivity to threat-predicting auditory cues, irrespective of the defensive strategy employed (*i.e*. freezing *vs* avoidance).

**Figure 5:**
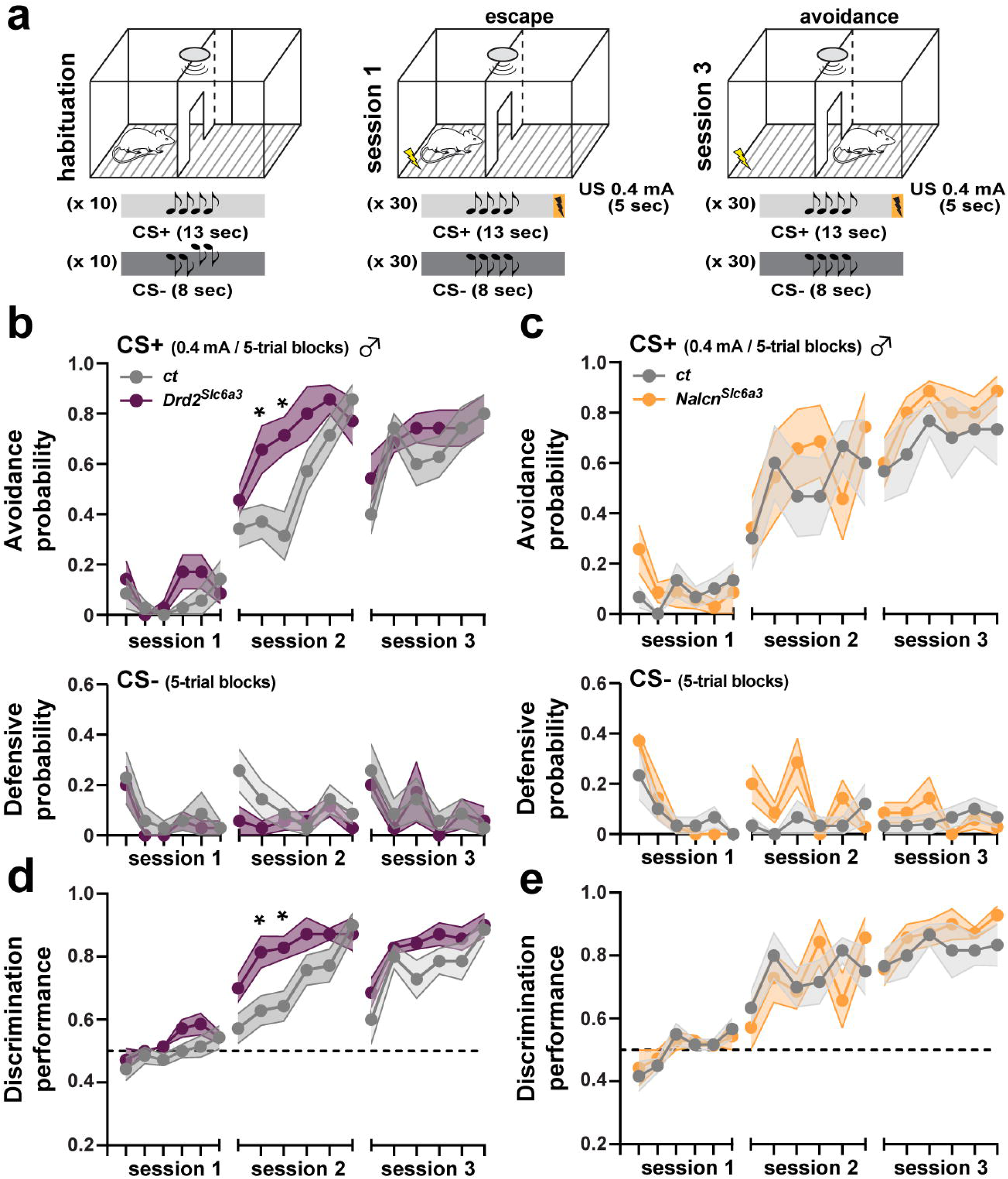
Discriminative active avoidance in male mice lacking *Drd2* or *Nalcn* in DA-neurons. (**a**) Schematic cartoon describing the protocol used to evaluate discriminative active avoidance learning. (**b**) Upper panel: avoidance probability in *ct* (grey) and *Drd2^Slc6a3^* (purple) mice representing the number of times the animal shuttled to the adjacent compartment during the CS+ presentation over 3 sessions (n = 7 *ct*, n = 7 *Drd2^Slc6a3^*). Lower panel: defensive probability representing the number of times the animal elicited an escape or avoidance response during the CS-presentation over 3 sessions. (**c**) Similar to (b) in *ct* (grey) and *Nalcn^Slc6a3^* (orange) (n = 6 *ct*, n = 7 *Nalcn^Slc6a3^*) (**d** and **e**) Discrimination performance across sessions between *ct* (grey) and *Drd2^Slc6a3^* (purple) mice in (**d**) and between *ct* (grey) and *Nalcn^Slc6a3^* (orange) mice in (**e**). Results are analyzed using two-way ANOVA repeated measures. *ct* vs *Drd2^Slc6a3^*, *p < 0.05, **p < 0.01. Statistical analysis detailed in Supplemental Table (5b-e).

## Discussion

Our study highlights the role of autoD2Rs in modulating the activity of VTA DA-neurons in response to aversive stimuli. By selectively deleting *Drd2* in midbrain DA-neurons, we observed significant enhancements in both excitatory and inhibitory responses of VTA DA-neurons to aversive stimuli. This suggests that autoD2Rs act as a regulatory brake, fine-tuning the encoding of both the value and saliency of negative stimuli. Furthermore, the lack of D2R in these neurons led to an enhanced ability to discriminate threat-predicting cues, particularly in male mice, suggesting a sex-specific regulatory role of autoD2R in threat processing. Finally, our results reveal that this modulation occurs independently of the pacemaker activity of DA-neurons or the coupling of D2R to NALCN channels, increasing our understanding of the molecular pathways by which D2R signaling modulates the activity of DA neurons and shapes behavioral responses to salient stimuli.

Midbrain DA-neurons are slow pacemakers that continuously fire action potentials, maintaining basal DA levels in their target regions. However, when exposed to salient stimuli, these neurons can switch to rapid burst firing, leading to transient DA increases (Grace et al., 2007; Schultz, 1998). DA-neurons excitability and DA release are tightly regulated by autoD2R which prevents excessive activation of DA-neurons and limits extracellular DA concentration (Ford, 2014; Wolf & Roth, 1990). The unaffected basal firing of VTA DA-neurons as well as the unchanged basal levels of extracellular DA in the nucleus accumbens in *Drd2^Slc6a3^* mice aligns with previous results obtained in D2R full or conditional knockout mice (Anzalone et al., 2012; Bello et al., 2011; Mercuri et al., 1997). While no differences in burst-like activity were found between *Drd2^Slc6a3^* and control mice, enhanced excitatory and inhibitory responses to salient aversive stimuli were observed. These results extend previous studies showing a heightened DA-neurons activity and DA release in *Drd2^Slc6a3^* mice in response to rewarding stimuli (Anzalone et al., 2012; Bello et al., 2011; Holroyd et al., 2015) indicating that autoD2Rs prevent exacerbated responses to both rewarding and aversive stimuli.

AutoD2Rs modulate the activity of DA-neurons through interactions with neighboring ion channels (Ford, 2014). While the coupling to GIRK2 channel has been extensively studied (Beaulieu & Gainetdinov, 2011; Ford, 2014; Neve et al., 2004), recent work suggests that the sodium leak channel NALCN is essential for sustaining the pacemaker activity of midbrain DA neurons (Cobb-Lewis et al., 2023; Philippart & Khaliq, 2018) and instrumental for scaling the D2R-mediated somatodendritic inhibition of DA neurons (Philippart & Khaliq, 2018). Our *in vivo* recordings extend these previous findings notably by demonstrating that NALCN conductance is crucial for the pacemaker activity of VTA DA-neurons (Cobb-Lewis et al., 2023; Philippart & Khaliq, 2018; Um et al., 2021). Although only one putative DA-neuron was recorded in *Nalcn^Slc6a3^* mice, the increased neuronal activity in response to salient aversive stimuli was preserved, suggesting that the enhanced responses observed in *Drd2^Slc6a3^* mice occurred independently of D2R-mediated NALCN inhibition. Further work is required to determine whether GIRK2 and/or βarrestin-2 signaling downstream autoD2R contributes to this regulation.

Previous studies have demonstrated that D2R signaling regulates locomotion. However, while the complete absence of D2R or specific invalidation in striatal medium spiny neurons leads to reduced movement (Baik et al., 1995; Kelly et al., 1998; Kharkwal et al., 2016; LeBlanc et al., 2020; Radl et al., 2018), specific deletion of either the long or short version of D2R does not affect locomotor activity (Radl et al., 2018; Usiello et al., 2000). Although increased locomotor activity has been associated with the lack of autoD2R signaling (Anzalone et al., 2012; Bello et al., 2011), no differences between *Drd2^Slc6a3^* and control male and female mice were found in the present study. Differences in how locomotor activity was measured and the experimental apparatus used likely accounted for this apparent discrepancy. Indeed, while initial studies used an open field, providing more opportunities for exploration, our study utilized a circular corridor in which habituation is rapid and exploration is reduced. Interestingly, locomotion remained unaltered in *Nalcn^Slc6a3^* mice. These results suggest that pacemaker activity and excitability of DA-neurons do not play a prominent role in the control of motor responses.

Increasing evidence suggests that midbrain DA transmission plays a critical role in processing threat-related events (Jo et al., 2018; Luo et al., 2018; Pezze & Feldon, 2004; Salinas-Hernández et al., 2018, 2023). Indeed, deficits in stimuli discrimination learning and generalized anxiety-like phenotypes have been observed in mice lacking the NR1 subunit of the NMDAR in midbrain DA neurons (Radke et al., 2019; Zweifel et al., 2011). D2R are particularly important for discriminative learning, as intact D2R signaling optimizes the differentiation between stimuli predicting safety or threat (Castell et al., 2024; De Bundel et al., 2016). Interestingly, our study revealed a sexually dimorphic effect of the lack of autoD2R on the reactivity to threat-predicting auditory cues. Only *Drd2^Slc6a3^* male mice displayed improved discrimination between auditory cues signaling (CS+) or not (CS-) threat, regardless of the defensive strategy employed (*i.e*. freezing *vs* avoidance). Interestingly, the phenotype observed in *Nalcn^Slc6a3^* male mice was radically different. First, discriminative active avoidance learning and sensitivity was not altered in *Nalcn^Slc6a3^* mice suggesting that engagement of dynamic defensive behavior (avoidance) can occurred independently of NALCN conductance and its subsequent role in the regulation of midbrain DA-neurons excitability. On the other hand, in discriminative auditory threat conditioning, *Nalcn^Slc6a3^* male mice failed to significantly discriminate between CS+ and CS-whatever the intensity shock used (0.6 or 0.7 mA), an impairment most likely resulting from the decreased sensitivity of DA-neurons encoding the value to aversive stimuli in *Nalcn^Slc6a3^* mice. Alternatively, future experiments will determine whether NALCN deletion in DA-neurons impairs animal’s ability to switch from one defensive behavioral strategy to another.

Finally, our study illustrates the need of considering sex as a biological variable when studying defensive behaviors. Indeed, evidence indicate that exploratory avoidance in response to threats is lower in female mice as compared to males (Castell et al., 2024; Yokota et al., 2017). Interestingly, extinction of conditioned threat in female mice is particularly sensitive to the hormonal status during the estrous cycle. In fact, females undergoing extinction training during proestrus perform similarly to males, whereas extinction is impaired during metestrus (Gruene et al., 2015; Lebrón-Milad et al., 2013; Milad et al., 2009; Velasco et al., 2019). Considering the existence of sex differences in the organization of the DA system and/or its modulation by sex steroid hormones (Di Paolo, 1994; Harrison & Tunbridge, 2008; Kritzer & Creutz, 2008; Riccardi et al., 2011; Vandegrift et al., 2017; Velasco et al., 2019), future studies will be necessary to determine whether the involvement of autoD2Rs in regulating defensive responses is influenced by the hormonal states.

In conclusion, our study unveils the importance of autoD2Rs signaling in serving as a brake to limit the excessive activation of VTA DA-neurons in response to salient aversive stimuli, thereby preventing exacerbated defensive responses evoked by threat predictive cues. This study also emphasizes the link between autoD2Rs dysfunctions and behavioral inflexibility, a feature common to several neuropsychiatric disorders characterized by maladaptive aversive behaviors and cognitive rigidity.

## Acknowledgments

The authors thank Pr Dejian Ren for generously providing the *Nalcn^f/f^* mice. We thank MRI platform (Biocampus). The authors also thank iExplore from IGF for their involvement in the maintenance and breeding of the colonies. This work was supported by CNRS, INSERM, Fondation pour la Recherche Médicale (EQU202203014705, EV), the French National Research Agency (ANR-20-CE14-0020; ANR-21-CE16-0028, EV), (ANR-19-CE16-0001, FM), (ANR-21-NEU2-0004-01, AM), the Labex “Ion Channel Science and Therapeutics” (ANR-11-LABX-0015, AM). E.P received funds from MINECO (RYC2020-029596-I) and MICINN (PID2021-125079OA-I00). G.G was supported by the Agence Nationale de la Recherche (ANR-21-CE14-0021-01; ANR-23-CE14-0014-02), Fédération pour la Recherche sur le Cerveau and Association France Parkinson, Institut Universtaire de France, and Université Paris Cité (Stratex_Idex-2023-028 EMERGENCE).

## Author contribution

L.C., A.G., F.M., and E.V. conceived the study and designed the experiments. L.C. and A.G. performed biochemical and behavioural experiments. A.C. and J.M.R. performed *in vivo* microdialysis experiments and analysis. E.P. performed Ribotag experiments. N. M. and G.G. performed and analysed western blot experiments. C.P.P. and R.N. performed and analyzed auditory brainstem responses. T.L.B., F.M. and P.F. designed, performed and analyzed *in vivo* juxtacellular recordings and *ex vivo* patch-clamp recordings. L.M. and C.N. performed the fluorescent *in situ* hybridization. C.N. and L.C. performed immunofluorescence experiments. A.B. provided insightful advices regarding the sex-specific effects. C. Bernat. performed genotyping. C. Bellone generated *Dat-Ribotag* transgenic mice. A.M. generated *Nalcn^Slc6a3^* transgenic mice. L.C., A.G., G.G., F.M. and E.V. wrote the manuscript the study with inputs from all authors.

## Declaration of interest

The authors declare no competing interests

## Notes

### Competing Interest Statement

The authors have declared no competing interest.

