## Supplemental material for "Midbrain dopamine D2R regulates the salience of threat-related events"

**Supplemental Figures**

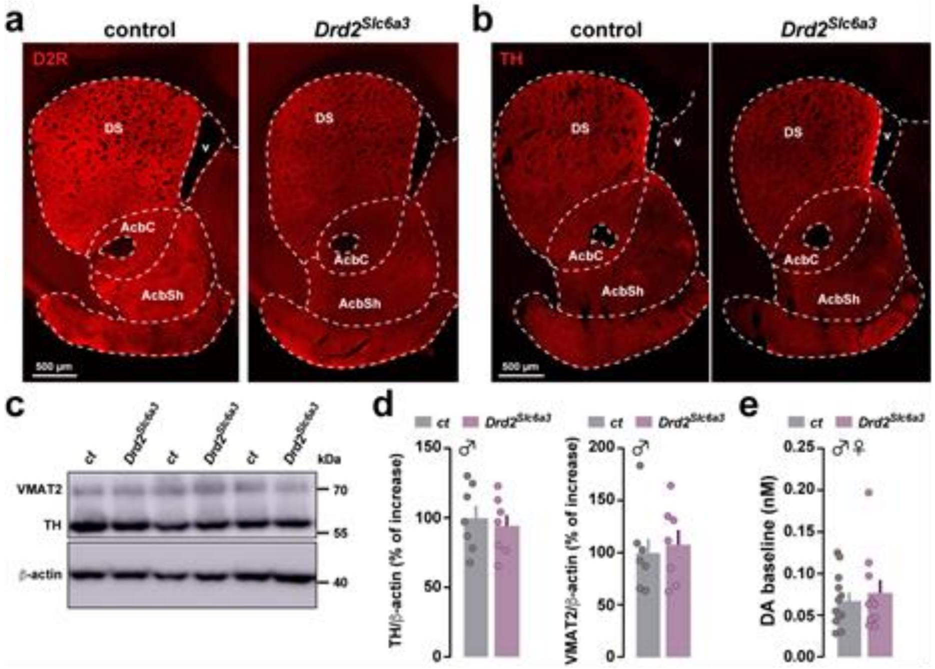

**Supplemental Figure 1: Expression of D2R, TH, VMAT2 and DA levels in control and *Drd2*$^{Slc6a3}$ mice. (a, b)** Coronal sections at the striatum level showing expression of D2R (a) and TH (b) in *ct* (left panel) and *Drd2*$^{Slc6a3}$ (right panel) mice. (**c, d**) Representative immunoblots (**c**) and quantifications of TH and VMAT2 expressions (**d**) in *ct* (n = 7) and *Drd2*$^{Slc6a3}$ (n = 7) mice. (e) Acb absolute basal levels of DA in dialysates collected from four consecutive fractions. Data correspond to the mean ± S.E.M. of the amount (nM) obtained in each experimental group. Statistical analysis detailed in Supplemental Table (S1d-e).

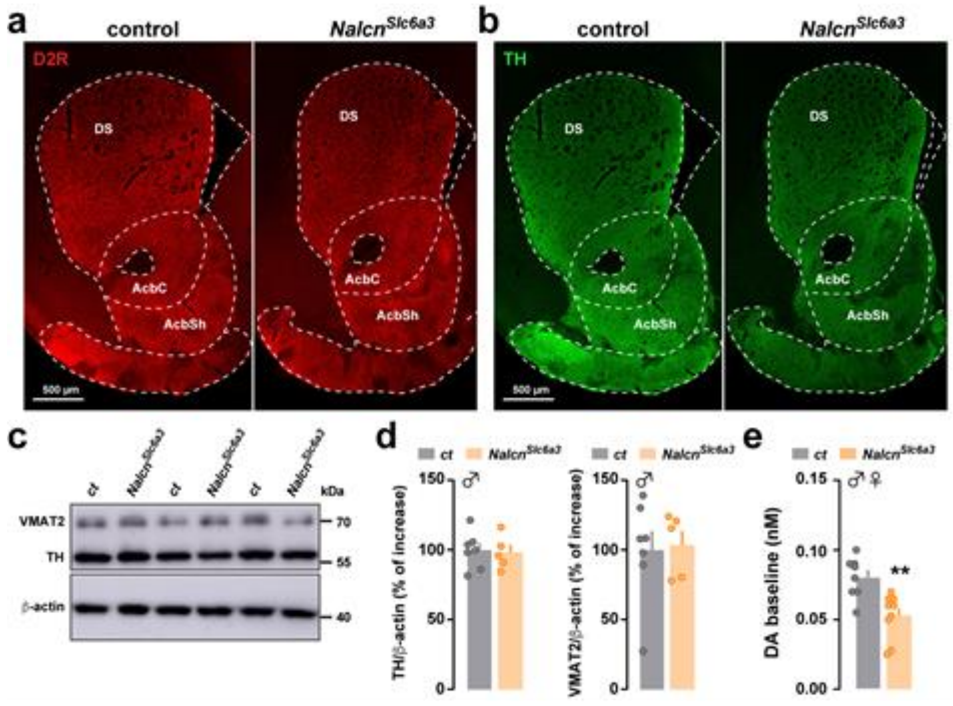

**Supplemental Figure 2: Expression of D2R, TH, VMAT2 and DA levels in control and *Nalcn*$^{Slc6a3}$ mice. (a, b)** Coronal sections at the striatum level showing expression of D2R (a) and TH (b) in *ct* (left panel) and *Nalcn*$^{Slc6a3}$ (right panel) mice. (**c, d**) Representative immunoblots (**c**) and quantifications of TH and VMAT2 expressions (**d**) in *ct* (n = 7) and *Nalcn*$^{Slc6a3}$ (n = 5) mice. (e) Acb absolute basal levels of DA in dialysates collected from four consecutive fractions. Data correspond to the mean ± S.E.M. of the amount (nM) obtained in each experimental group. Statistical analysis detailed in Supplemental Table (S2d-e).

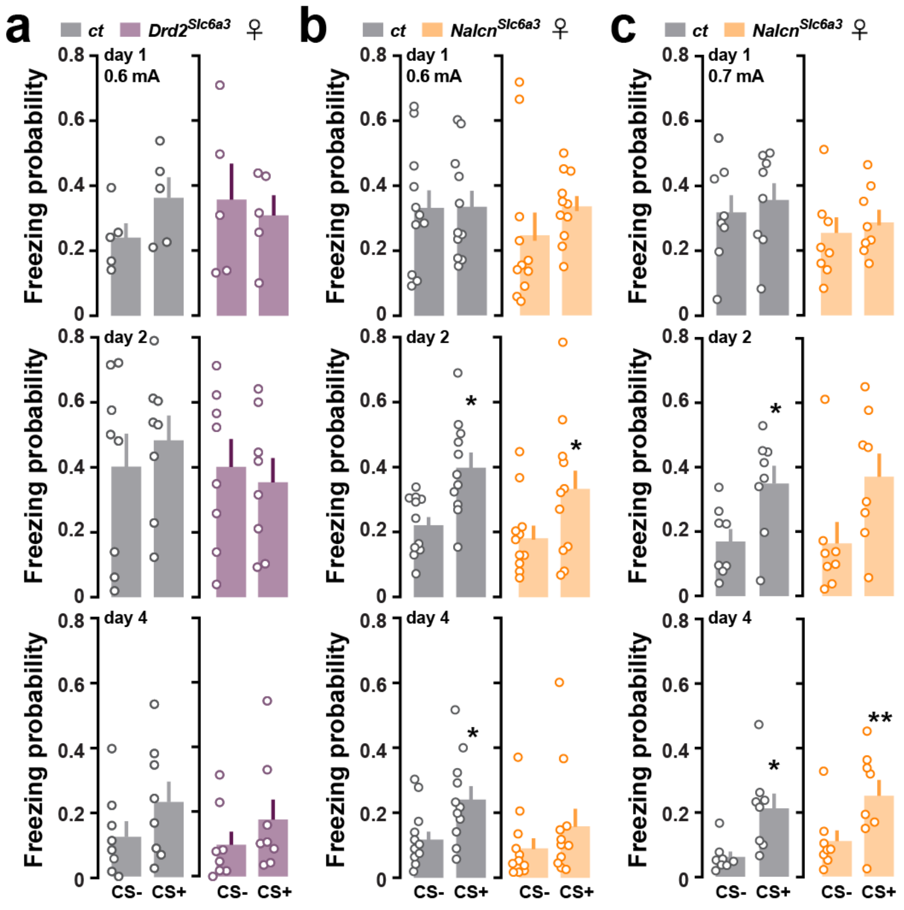

**Supplemental Figure 3: Discriminative learning to threat-predicting cues in female mice lacking *Drd2* or *Nalcn* in DA-neurons**. (**a**) Probability of CS- and CS+ to evoke freezing responses in *ct* (grey) and *Drd2*$^{Slc6a3}$ (purple) mice during the conditioning (day 1, upper panel), the test (day 2, middle panel) and after extinction (day 4, lower panel) (n = 8 *ct*, n = 8 *Drd2*$^{Slc6a3}$). (**b**) Similar to (**a**) in *ct* (grey) and *Nalcn*$^{Slc6a3}$ (orange) mice (n = 11 *ct*, n = 11 *Nalcn*$^{Slc6a3}$). (**c**) Similar to (**a**) in *ct* (grey) and *Nalcn*$^{Slc6a3}$ (orange) mice exposed to a 0.7mA shock (n = 8 *ct*, n = 8 *Nalcn*$^{Slc6a3}$). Results are analyzed using paired *t*-test. CS- vs CS+, *$p < 0.05$, **$p < 0.01$. Statistical analysis detailed in Supplemental Table (S3a-c).

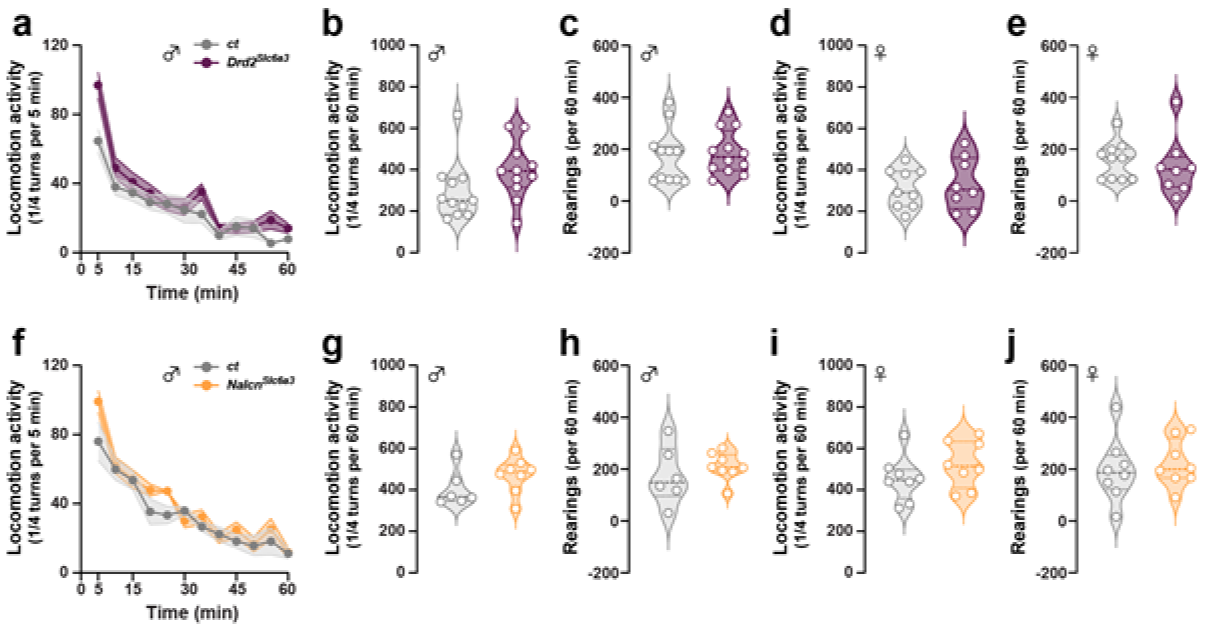

**Supplemental Figure 4: Locomotion and rearing behavior in male and female mice lacking *Drd2* or *Nalcn* in DA-neurons.** (**a, b, d**) Spontaneous locomotor activity in males (**a,b**) and females (**d**) *ct* (grey) and *Drd2*$^{Slc6a3}$ (purple) mice over 60 min in a circular corridor (males: n = 11 *ct*, n = 11 *Drd2*$^{Slc6a3}$ , females: n = 9 *ct*, n = 8 *Drd2*$^{Slc6a3}$). (**c, e**) Vertical activity (rearings) over 60 min in males (**c**) and females (**e**) in a circular corridor (males: n = 11 *ct*, n = 12 *Drd2*$^{Slc6a3}$, females: n = 10 *ct*, n = 8 *Drd2*$^{Slc6a3}$). (f**, g, i**) Spontaneous locomotor activity in males (**f, g**) and females (**i**) *ct* (grey) and *Nalcn*$^{Slc6a3}$ (orange) mice over 60 min in a circular corridor (males: n = 6 *ct*, n = 8 *Nalcn*$^{Slc6a3}$, females: n = 8 *ct*, n = 9 *Nalcn*$^{Slc6a3}$). (**h, j**) Vertical activity (rearings) over 60 min in males (**h**) and females (**j**) in a circular corridor (males: n = 6 *ct*, n = 8 *Nalcn*$^{Slc6a3}$, females: n = 8 *ct*, n = 8 *Nalcn*$^{Slc6a3}$). Statistical analysis detailed in Supplemental Table (S4a-j).

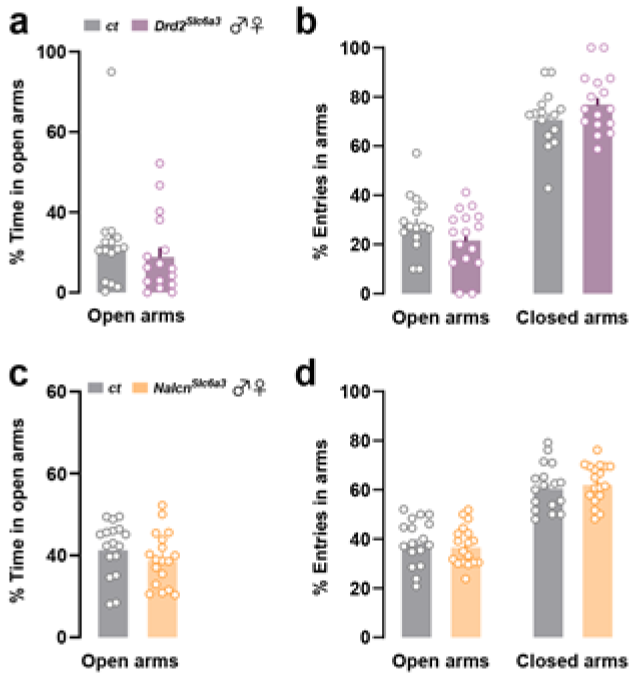

**Supplemental Figure 5: Anxiety-like behavior in male and female mice lacking *Drd2* or *Nalcn* in DA-neurons.** (**a, b**) Percentage of time spent open arms (**a**) and entries in open and closed arms (**b**) during 5 minutes in the elevated plus maze in males and females *ct* (grey) and $Drd2^{Slc6a3}$ (purple) mice (n = 15 *ct*, n = 16 $Drd2^{Slc6a3}$). (**c, d**) Percentage of of time spent open arms (**c**) and entries in open and closed arms (**d**) during 5 minutes in the elevated plus maze in males and females *ct* (grey) and $Nalcn^{Slc6a3}$ (orange) mice (n = 17 *ct*, n = 18 $Nalcn^{Slc6a3}$). Statistical analysis detailed in Supplemental Table (S5a-d).

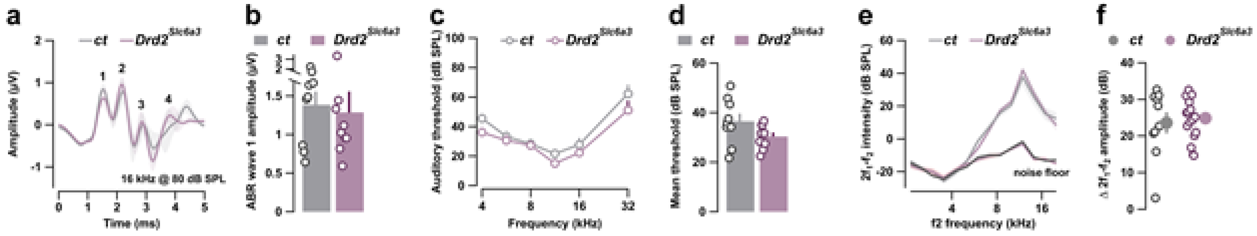

**Supplemental Figure 6: Normal hearing in *Drd2$^{Slc6a3}$* mice.** (**a**, **b**) Mean auditory brainstem response (ABR) ± SEM recordings evoked by 16 kHz tone burst at 80 dB SPL (**a**) and mean wave 1 ABR ± SEM amplitude (**b**) from control and *Drd2$^{Slc6a3}$* mice (n = 9 cochlea from 9 mice in *ct*, n = 9 cochlea from 9 mice in *Drd2$^{Slc6a3}$*). Numbers in (**a**) indicate the synchronous activation of nuclei along the ascending auditory pathway: (1) spiral ganglion neuron, (2) cochlear nuclei, (3) superior olive and (4) lateral lemniscus. (**c**, **d**) Mean ABR ± SEM audiograms (**c**) and mean auditory threshold (**d**) from control and *Drd2$^{Slc6a3}$* mice (n = 9 cochlea from 9 mice in *ct*, n = 9 cochlea from 9 mice in *Drd2$^{Slc6a3}$*). (**e**) Distortion product otoacoustic emissions from control and *Drd2$^{Slc6a3}$* mice (n = 10 cochlea from 5 mice in *ct*, n = 16 cochlea from 8 mice in *Drd2$^{Slc6a3}$*). The $2f_1$-$f_2$ amplitude level, reflecting the amplification mechanism within the cochlea, is shown as a function of $f_2$ frequency. The black lines indicate the background noise level. (**f**) Mean $2f_1$-$f_2$ ± SEM amplitude level from (**e**) measured between 5 and 20 kHz. Statistical analysis detailed in Supplemental Table (S6a-b).

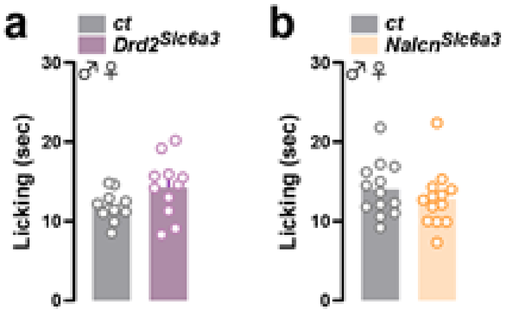

**Supplemental Figure 7: Nociceptive threshold of mice lacking *Drd2* or *Nalcn* in DA-neurons.** (**a, b**) Latency to the first licking in the hotplate test in *ct* and *Drd2$^{Slc6a3}$* mice (**a**) (n = 12 *ct*, n = 11 *Drd2$^{Slc6a3}$*) and *ct* and *Nalcn$^{Slc6a3}$* mice (**b**) (n = 13 *ct*, n = 13 *Nalcn$^{Slc6a3}$*). Statistical analysis detailed in Supplemental Table (S7a-b).

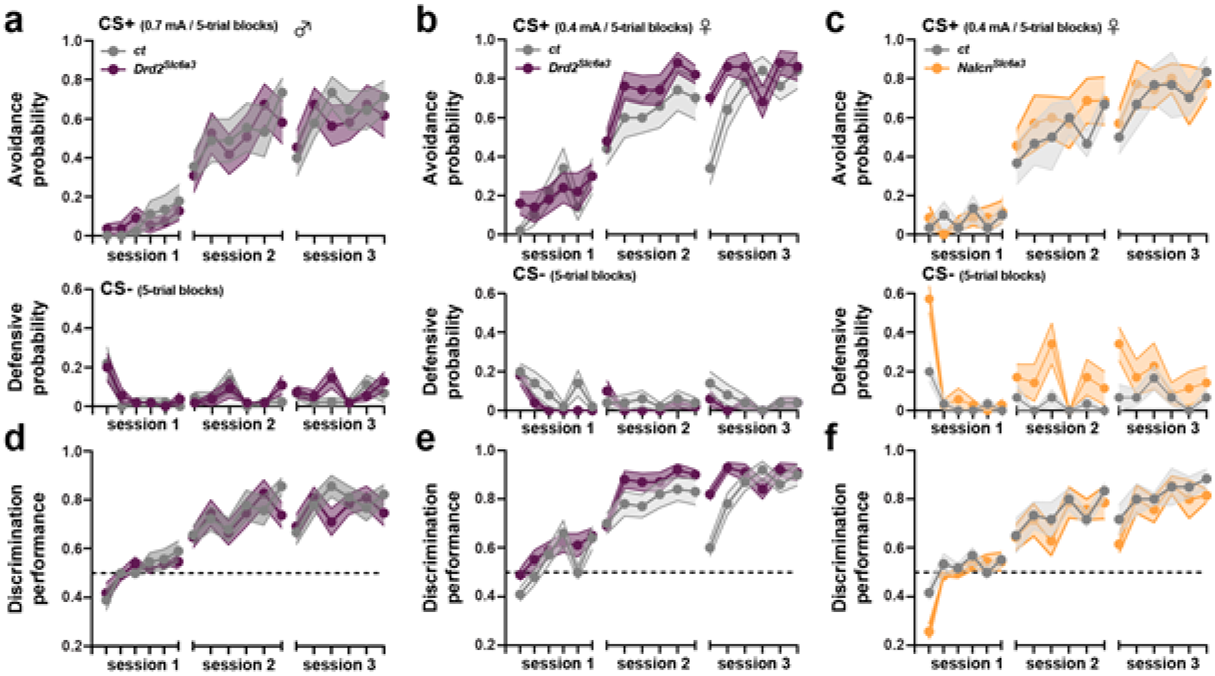

**Supplemental Figure 8: Active avoidance in male and female mice lacking *Drd2* or *Nalcn* in DA-neurons**. (**a, b**) Avoidance probability (number of times the animal shuttled to the adjacent compartment during the CS+, upper panel) and defensive probability (number of times the animal shuttled to the adjacent compartment during the CS-, lower panel) among 3 sessions in male (a) and female (b) *ct* and $Drd2^{Slc6a3}$ mice. (**c**) Similar to (a), avoidance and defensive probability for female *ct* and $Nalcn^{Slc6a3}$ mice. (**d, e**) Discrimination performance in male (d) and female (e) *ct* and $Drd2^{Slc6a3}$ mice (males: n = 9 *ct*, n = 11 $Drd2^{Slc6a3}$, females: n = 10 *ct*, n = 10 $Drd2^{Slc6a3}$). (**f**) Discrimination performance in female *ct* and $Nalcn^{Slc6a3}$ mice (n = 6 *ct*, n = 7 $Nalcn^{Slc6a3}$). Statistical analysis detailed in Supplemental Table (S8a-f).

**Supplemental Table 1**

List of primers

| Gene | Primer Forward | Primer Reverse |
|---|---|---|
| *Aif1* | CCCCCAGCCAAGAAAGCTAT | GCCCCACCGTGTGACATC |
| *Cnp* | GCTGCACTGTACAACCAAATTCTG | ACCTCCTGCTGGGCGTATT |
| *Drd2* | CTCTTTGGACTCAACAACACAGA | AAGGGCACGTAGAACGAGAC |
| *Gad1* | TTGTGCTTTGCTGTGTTTTAGAGA | CCCCCTGCCCAAAGATAGAC |
| *Gapdh* | CTCAACTACATGGTCTACATGTTCCA | CCCATTCTCGGCCTTGACT |
| *Gfap* | AGCGAGCGTGCAGAGATGA | AGGAAGCGGACCTTCTCGAT |
| *Slc6a3* | AGCATTGTGACCTTCAGACCCC | AGCTGGCGGTCTTTCTCAGG |
| *Th* | CCGTGCAGCCCTACCAAGAT | CCGGATGGTGTGAGGACTGTC |

**Supplemental Table 2**

| Figure | Measure | | Groups | Statistical Analysis |
|---|---|---|---|---|
| 2a | Cumulative probability of spontaneous frequency and % of SWB | male | Control (neurons n = 107)<br>*Drd2*$^{Slc6a3}$ (neurons n = 115) | **Kolmogorov-Smirnov test**<br>Frequency<br>D = 0.14, p = 0.2295<br>% SWB<br>D = 0.098, p = 0.6625 |
| 2b | Firing variation in pinch-activated DA neurons | male | Control (neurons n = 45)<br>*Drd2*$^{Slc6a3}$ (neurons n = 33) | **Wilcoxon test**<br>Mean: W = 618, p = 0.21<br>Max: W = 945, p = 0.04* |
| 2c | Firing variation in pinch-inhibited DA neurons | male | Control (neurons n = 45)<br>*Drd2*$^{Slc6a3}$ (neurons n = 49) | **Wilcoxon test**<br>Mean: W = 1336, p = 0.047*<br>Max: W = 912, p = 0.15 |
| 3d | Spontaneous activity<br><br>Cumulative probability | male | Control (n = 14 neurons, n = 2 mice)<br>*Nalcn*$^{Slc6a3}$ (n = 22 neurons, n = 3 mice) | **Wilcoxon test**<br>W = 212, p = 0.04608*<br><br>**Kolmogorov-Smirnov test**<br>D = 0.461, p = 0.05266 |
| 3e | Resting membrane potential | male | Control (n = 14 neurons, n = 2 mice)<br>*Nalcn*$^{Slc6a3}$ (n = 22 neurons, n = 3 mice) | **Kolmogorov-Smirnov test**<br>D = 0.47403, p = 0.02769* |
| 3f | Current response curve | male | Control (n = 14 neurons, n = 2 mice)<br>*Nalcn*$^{Slc6a3}$ (n = 22 neurons, n = 3 mice) | **Two-way ANOVA repeated measures**<br>Performed on all currents:<br>Genotype<br>$F_{(1, 33)}$ = 4.057, p = 0.0522<br>Courant<br>$F_{(10, 330)}$ = 30.631, p<2e-16***<br>Interaction<br>$F_{(10, 330)}$ = 1.518, p = 0.131<br><br>Performed only on small currents (0, 10, 20, 30 pA);<br>Genotype<br>$F_{(1, 33)}$ = 6.691, p = 0.0143*<br>Courant<br>$F_{(3, 99)}$ = 26.048, p = 1.67e-12***<br>Interaction<br>$F_{(3, 99)}$ = 3.491, p = 0.0185*<br><br>**Wilcoxon test**<br>Control vs. *Nalcn*$^{Slc6a3}$<br>0 pA: W = 165, p = 0.06801<br>10 pA: W = 205.5, p = 0.01517* |

| | | | | |
|---|---|---|---|---|
| | | | | 20 pA: W = 208, p = 0.01693*<br>30 pA: W = 202, p = 0.03186* |
| 3g | I/V relationship | male | Control (n = 14 neurons, n = 2 mice)<br>*Nalcn*$^{Slc6a3}$ (n = 22 neurons, n = 3 mice) | **Two-way ANOVA repeated measures**<br>Genotype<br>$F_{(1, 34)} = 0.029$, $p = 0.866$<br>Courant<br>$F_{(14,476)} = 381.103$, $p<2e-16$***<br>Interaction<br>$F_{(14,476)} = 0.016$, $p = 1$ |
| 3h | Ih current | male | Control (n = 14 neurons, n = 2 mice)<br>*Nalcn*$^{Slc6a3}$ (n = 20 neurons, n = 3 mice) | **Wilcoxon test**<br>W = 178, p = 0.1922 |
| 3i | Spontaneous EPSC Amplitude<br><br>Cumulative probability<br><br><br>Frequency<br><br><br>Cumulative probability | male | Control (n = 13 neurons, n = 2 mice)<br>*Nalcn*$^{Slc6a3}$ (n = 19 neurons, n = 3 mice) | **Welch two-sample t-test**<br>$t_{19.629} = 2.7604$, $p = 0.01221$*<br><br>**Kolmogorov-Smirnov test**<br>D = 0.41902, p < 2.2e-16***<br><br>**Wilcoxon test**<br>W = 61, p = 0.01577*<br><br>**Kolmogorov-Smirnov test**<br>D = 0.27887, p < 2.2e-16* |
| 3k | Mean number of spike | male | *Nalcn*$^{Slc6a3}$ (n = 5 neurons, n = 2 mice) | **Wilcoxon test**<br>W = 0, p = 0.021*<br>W = 1, p = 0.02* |
| 4b | Freezing Probability 0.6 mA Conditioning (day1, upper)<br><br><br>Test (day 2, middle)<br><br><br>Extinction (day 4, lower) | male | <br>Control (mice n = 10)<br>*Drd2*$^{Slc6a3}$ (mice n = 12)<br><br>Control (mice n = 10)<br>*Drd2*$^{Slc6a3}$ (mice n = 12)<br><br>Control (mice n = 11)<br>*Drd2*$^{Slc6a3}$ (mice n = 12) | **Paired t-test**<br>$t_9 = 0.996$, $p = 0.3451$<br>$t_{11} = 2.764$, $p = 0.0184$*<br><br>$t_9 = 7.597$, $p < 0.0001$***<br>$t_{11} = 6.283$, $p < 0.0001$***<br><br>$t_9 = 1.945$, $p = 0.836$<br>$t_{11} = 3.743$, $p = 0.0032$** |
| 4c | Freezing Probability 0.6 mA Conditioning (day1, upper)<br><br><br>Test (day 2, middle)<br><br><br>Extinction (day 4, lower) | male | <br>Control (mice n = 7)<br>*Nalcn*$^{Slc6a3}$ (mice n = 7)<br><br>Control (mice n = 7)<br>*Nalcn*$^{Slc6a3}$ (mice n = 7)<br><br>Control (mice n = 7)<br>*Nalcn*$^{Slc6a3}$ (mice n = 7) | **Paired t-test**<br>$t_6 = 0.43$, $p = 0.6822$<br>$t_6 = 1.558$, $p = 0.1702$<br><br>$t_6 = 0.1159$, $p = 0.9115$<br>$t_6 = 1.877$, $p = 0.1095$<br><br>$t_6 = 1.582$, $p = 0.1647$<br>$t_6 = 1.658$, $p = 0.1485$ |
| 4d | Freezing Probability 0.7 mA | male | | **Paired t-test** |

| | | | | |
|---|---|---|---|---|
| | Conditioning (day1, upper)<br><br><br>Test (day 2, middle)<br><br><br>Extinction (day 4, lower) | | Control (mice n = 8)<br>*Nalcn*$^{Slc6a3}$ (mice n = 9)<br><br>Control (mice n = 8)<br>*Nalcn*$^{Slc6a3}$ (mice n = 9)<br><br>Control (mice n = 8)<br>*Nalcn*$^{Slc6a3}$ (mice n = 9) | $t_7$ = 1.706, p = 0.1318<br>$t_8$ = 1.134, p = 0.2894<br><br>$t_7$ = 3.419, p = 0.0112*<br>$t_8$ = 2.128, p = 0.0660<br><br>$t_7$ = 4.267, p = 0.0037**<br>$t_8$ = 1.640, p = 0.1396 |
| 5b | Avoidance probability<br><br><br><br>Defensive probability | male | Control (mice n = 7)<br>*Drd2*$^{Slc6a3}$ (mice n = 7) | **Two-way ANOVA repeated measures**<br>Genotype<br>$F_{(1, 12)}$ = 6.507 , p = 0.0254*<br>$F_{(1, 12)}$ = 2.321 , p = 0.1536 |
| 5c | Avoidance probability<br><br><br><br>Defensive probability | male | Control (mice n = 6)<br>*Nalcn*$^{Slc6a3}$ (mice n = 7) | **Two-way ANOVA repeated measures**<br>Genotype<br>$F_{(1, 11)}$ = 0.4234 , p = 0.5286<br>$F_{(1, 11)}$ = 2.323 , p = 0.1517 |
| 5d | Discrimination performance | male | Control (mice n = 7)<br>*Drd2*$^{Slc6a3}$ (mice n = 7) | **Two-way ANOVA repeated measures**<br>Genotype<br>$F_{(1, 12)}$ = 8.823 , p = 0.0117* |
| 5e | Discrimination performance | male | Control (mice n = 6)<br>*Nalcn*$^{Slc6a3}$ (mice n = 7) | **Two-way ANOVA repeated measures**<br>Genotype<br>$F_{(1, 11)}$ = 0.09917 , p = 0.7587 |
| S1d | TH<br><br><br>VMAT2 | male | Control (mice n = 7)<br>*Drd2*$^{Slc6a3}$ (mice n = 7)<br><br>Control (mice n = 7)<br>*Drd2*$^{Slc6a3}$ (mice n = 7) | **Unpaired t-test**<br>$t_{12}$ = 0.4876, p = 0.6346<br><br>$t_{12}$ = 0.3689, p = 0.7186 |
| S1e | DA levels | male and female | Control (mice n = 10)<br>*Drd2*$^{Slc6a3}$ (mice n = 12) | **Unpaired t-test**<br>$t_{20}$ = 0.4807, p = 0.6360 |
| S2d | TH<br><br><br>VMAT2 | male | Control (mice n = 7)<br>*Nalcn*$^{Slc6a3}$ (mice n = 5)<br><br>Control (mice n = 7)<br>*Nalcn2*$^{Slc6a3}$ (mice n = 5) | **Unpaired t-test**<br>$t_{10}$ = 0.2095, p = 0.8383<br><br>$t_{10}$ = 0.1988, p = 0.8464 |
| S2e | DA levels | male and female | Control (mice n = 8)<br>*Nalcn2*$^{Slc6a3}$ (mice n = 9) | **Unpaired t-test**<br>$t_{15}$ = 3.661, p = 0.0023** |
| S3a | Freezing Probability 0.6 mA<br>Conditioning (day1, upper)<br><br><br>Test (day 2, middle) | female | <br>Control (mice n = 5)<br>*Drd2*$^{Slc6a3}$ (mice n = 5)<br><br>Control (mice n = 8)<br>*Drd2*$^{Slc6a3}$ (mice n = 8) | **Paired t-test**<br>$t_4$ = 2.424, p = 0.0724<br>$t_4$ = 0.3700, p = 0.7266<br><br>$t_7$ = 1.353, p = 0.2182<br>$t_7$ = 0.3367, p = 0.7462 |

| | | | | |
|---|---|---|---|---|
| | Extinction (day 4, lower) | | Control (mice n = 8)<br>*Drd2*$^{Slc6a3}$ (mice n = 8) | $t_7$ = 1.578, p = 0.1586<br>$t_7$ = 2.195, p = 0.1852 |
| S3b | Freezing Probability 0.6 mA<br>Conditioning (day1, upper)<br><br><br>Test (day 2, middle)<br><br><br>Extinction (day 4, lower) | female | <br>Control (mice n = 11)<br>*Nalcn*$^{Slc6a3}$ (mice n = 11)<br><br>Control (mice n = 11)<br>*Nalcn*$^{Slc6a3}$ (mice n = 11)<br><br>Control (mice n = 11)<br>*Nalcn*$^{Slc6a3}$ (mice n = 11) | **Paired t-test**<br>$t_4$ = 0.02481, p = 0.9807<br>$t_4$ = 1.342, p = 0.2093<br><br>$t_{10}$ = 2.452, p = 0.0342*<br>$t_{10}$ = 2.334, p = 0.0417*<br><br>$t_{10}$ = 2.873, p = 0.0166*<br>$t_{10}$ = 2.129, p = 0.0591 |
| S3c | Freezing Probability 0.7 mA<br>Conditioning (day1, upper)<br><br><br>Test (day 2, middle)<br><br><br>Extinction (day 4, lower) | female | <br>Control (mice n = 8)<br>*Nalcn*$^{Slc6a3}$ (mice n = 8)<br><br>Control (mice n = 8)<br>*Nalcn*$^{Slc6a3}$ (mice n = 8)<br><br>Control (mice n = 8)<br>*Nalcn*$^{Slc6a3}$ (mice n = 8) | **Paired t-test**<br>$t_7$ = 0.7403, p = 0.4889<br>$t_7$ = 1.285, p = 0.239<br><br>$t_7$ = 2.482, p = 0.0421*<br>$t_7$ = 2.024, p = 0.0827<br><br>$t_7$ = 2.775, p = 0.0275*<br>$t_7$ = 4.527, p = 0.0027** |
| S4a | Locomotion | male | Control (mice n = 9)<br>*Drd2*$^{Slc6a3}$ (mice n = 8) | **Two-way ANOVA repeated measures**<br>Genotype<br>$F_{(1, 20)}$ = 0.8483, p = 0.3680 |
| S4b | Locomotion | male | Control (mice n = 11)<br>*Drd2*$^{Slc6a3}$ (mice n = 11) | **Unpaired t-test**<br>$t_{20}$ = 0.9210, p = 0.3680 |
| S4c | Rearings | male | Control (mice n = 11)<br>*Drd2*$^{Slc6a3}$ (mice n = 12) | **Unpaired t-test**<br>$t_{21}$ = 0.3002, p = 0.7670 |
| S4d | Locomotion | female | Control (mice n = 9)<br>*Drd2*$^{Slc6a3}$ (mice n = 8) | **Unpaired t-test**<br>$t_{15}$ = 0.4183, p = 0.6816 |
| S4e | Rearings | female | Control (mice n = 10)<br>*Drd2*$^{Slc6a3}$ (mice n = 8) | **Unpaired t-test**<br>$t_{16}$ = 0.5060, p = 0.6198 |
| S4f | Locomotion | male | Control (mice n = 6)<br>*Nalcn*$^{Slc6a3}$ (mice n = 8) | **Two-way ANOVA repeated measures**<br>Genotype<br>$F_{(1, 12)}$ = 2.108, p = 0.1722 |
| S4g | Locomotion | male | Control (mice n = 6)<br>*Nalcn*$^{Slc6a3}$ (mice n = 8) | **Unpaired t-test**<br>$t_{12}$ = 1.452, p = 0.1722 |
| S4h | Rearings | male | Control (mice n = 6)<br>*Nalcn*$^{Slc6a3}$ (mice n = 8) | **Unpaired t-test**<br>$t_{12}$ = 1.021, p = 0.3275 |
| S4i | Locomotion | female | Control (mice n = 8)<br>*Nalcn*$^{Slc6a3}$ (mice n = 8) | **Unpaired t-test**<br>$t_{14}$ = 1.295, p = 0.2163 |
| S4j | Rearings | female | Control (mice n = 8)<br>*Nalcn*$^{Slc6a3}$ (mice n = 8) | **Unpaired t-test**<br>$t_{14}$ = 0.4702, p = 0.6455 |
| S5a | Time in open arms | male and female | Control (mice n = 15)<br>*Drd2*$^{Slc6a3}$ (mice n = 16) | **Unpaired t-test**<br>$t_{29}$ = 0.7115, p = 0.4825 |
| S5b | Entries in arms | male and female | Control (mice n = 15) | **Two-way ANOVA** |

| | | | | |
|---|---|---|---|---|
| | | | *Drd2*$^{Slc6a3}$ (mice n = 16) | **Mixed model**<br>Genotype<br>$F_{(1, 66)} = 0.000$ , $p = 0.999$ |
| S5c | Time in open arms | male and female | Control (mice n = 17)<br>*Nalcn*$^{Slc6a3}$ (mice n = 18) | **Unpaired t-test**<br>$t_{33} = 1.160$, $p = 0.2542$ |
| S5d | Entries in arms | male and female | Control (mice n = 17)<br>*Nalcn*$^{Slc6a3}$ (mice n = 18) | **Two-way ANOVA**<br>**Mixed model**<br>Genotype<br>$F_{(1, 58)} = 0.000$ , $p < 0.999$ |
| S6b | Mean wave 1 ABR amplitude | male | Control (mice n = 9, cochlea n = 9)<br>*Drd2*$^{Slc6a3}$ (mice n = 9, cochlea n = 9) | **Mann–Whitney Wilcoxon's test**<br>$p = 0.3098$ |
| S6d | mean auditory threshold | male | Control (mice n = 9, cochlea n = 9)<br>*Drd2*$^{Slc6a3}$ (mice n = 9, cochlea n = 9) | **Mann–Whitney Wilcoxon's test**<br>$p = 0.132719$ |
| S6f | mean $2f_1$-$f_2$ amplitude | male | Control (mice n = 5, cochlea n = 10)<br>*Drd2*$^{Slc6a3}$ (mice n = 8, cochlea n=16) | **Mann–Whitney Wilcoxon's test**<br>$p = 0.833029$ |
| S7a | Licking latency | male and female | Control (mice n = 12)<br>*Drd2*$^{Slc6a3}$ (mice n = 11) | **Unpaired t-test**<br>$t_{21} = 2.035$, $p = 0.0546$ |
| S7b | Licking latency | male and female | Control (mice n = 13)<br>*Nalcn*$^{Slc6a3}$ (mice n = 13) | **Unpaired t-test**<br>$t_{24} = 0.8938$, $p = 0.3803$ |
| S8a | Avoidance probability<br><br><br>Defensive probability | male | Control (mice n = 9)<br>*Drd2*$^{Slc6a3}$ (mice n = 11) | **Two-way ANOVA repeated measures**<br>Genotype<br>$F_{(1, 18)} = 0.04825$ , $p = 0.8286$<br>$F_{(1, 18)} = 0.4615$ , $p = 0.5056$ |
| S8b | Avoidance probability<br><br><br><br>Defensive probability | female | Control (mice n = 10)<br>*Drd2*$^{Slc6a3}$ (mice n = 10) | **Two-way ANOVA repeated measures**<br>Genotype<br>$F_{(1, 18)} = 1.289$ , $p = 0.2711$<br>$F_{(1, 18)} = 2.507$ , $p = 0.1307$ |
| S8c | Avoidance probability<br><br><br><br>Defensive probability | female | Control (mice n = 6)<br>*Nalcn*$^{Slc6a3}$ (mice n = 7) | **Two-way ANOVA repeated measures**<br>Genotype<br>$F_{(1, 11)} = 0.2292$ , $p = 0.6415$<br>$F_{(1, 11)} = 5.678$ , $p = 0.0363$* |
| S8d | Discrimination performance | male | Control (mice n = 9)<br>*Drd2*$^{Slc6a3}$ (mice n = 11) | **Two-way ANOVA repeated measures**<br>Genotype<br>$F_{(1, 18)} = 0.1541$ , $p = 0.6992$ |
| S8e | Discrimination performance | female | Control (mice n = 10)<br>*Drd2*$^{Slc6a3}$ (mice n = 10) | **Two-way ANOVA repeated measures**<br>Genotype |

| | | | | |
|---|---|---|---|---|
| | | | | $F_{(1, 18)}$ = 4.203 , p = 0.0552 |
| S8f | Discrimination performance | female | Control (mice n = 6)<br>*Nalcn*$^{Slc6a3}$ (mice n = 7) | **Two-way ANOVA repeated measures**<br>Genotype<br>$F_{(1, 11)}$ = 0.9456 , p = 0.3517 |
